# A dual role for H2A.Z.1 in modulating the dynamics of RNA Polymerase II initiation and elongation

**DOI:** 10.1101/2020.06.22.165373

**Authors:** Constantine Mylonas, Alexander L. Auld, Choongman Lee, Ibrahim I. Cisse, Laurie A. Boyer

## Abstract

RNAPII pausing immediately downstream of the transcription start site (TSS) is a critical rate limiting step at most metazoan genes that allows fine-tuning of gene expression in response to diverse signals^1–5^. During pause-release, RNA Polymerase II (RNAPII) encounters an H2A.Z.1 nucleosome^6–8^, yet how this variant contributes to transcription is poorly understood. Here, we use high resolution genomic approaches^2,9^ (NET-seq and ChIP-nexus) along with live cell super-resolution microscopy (tcPALM)^10^ to investigate the role of H2A.Z.1 on RNAPII dynamics in embryonic stem cells (ESCs). Using a rapid, inducible protein degron system^11^ combined with transcriptional initiation and elongation inhibitors, our quantitative analysis shows that H2A.Z.1 slows the release of RNAPII, impacting both RNAPII and NELF dynamics at a single molecule level. We also find that H2A.Z.1 loss has a dramatic impact on nascent transcription at stably paused, signal-dependent genes. Furthermore, we demonstrate that H2A.Z.1 inhibits re-assembly and re-initiation of the PIC to reinforce the paused state and acts as a strong additional pause signal at stably paused genes. Together, our study suggests that H2A.Z.1 fine-tunes gene expression by regulating RNAPII kinetics in mammalian cells.

## Introduction

Eukaryotic transcription is a highly controlled process facilitated by a compendium of protein complexes that regulate RNA Polymerase II (RNAPII) recruitment as well as transcription initiation, elongation and termination ultimately culminating in genic output^3–5^. In metazoans, RNAPII promoter-proximal pausing is now widely recognized as a key rate-limiting step that occurs between +25 to +50 nucleotides downstream of the transcription start site (TSS) and prior to the first nucleosome^1–3^. Current models suggest that RNAPII pausing provides a critical window of opportunity to fine tune gene expression in response to signaling cues^12^. A diverse set of positive and negative factors regulating RNAPII pausing have been extensively studied both *in vitro*^13–15^ and *in vivo*^3,16^, yet how promoter nucleosomes affect pause-release remains poorly understood. At most genes in eukaryotes, the highly conserved histone H2A variant H2A.Z marks nucleosomes that flank the nucleosome depleted region (NDR) around the TSS^7,17–19^. H2A.Z is essential for early metazoan development^20,21^ and is necessary for proper induction of differentiation programs^19,22–24^. Conflicting reports suggest both positive and negative functions for H2A.Z in gene regulation. For example, work in *Drosophila* showed that H2A.Z (H2avD) lowers the energy barrier for RNAPII passage^7^, whereas recent biophysical studies using optical tweezers on *in vitro* reconstituted templates demonstrated that mammalian H2A.Z nucleosomes are more stable and pose a greater barrier to RNAPII compared to canonical histone H2A^6^. Notably, disruption of proper H2A.Z incorporation contributes to a growing list of human diseases including cancer^23,25–27^ underscoring the need for dissecting its mechanistic link to transcription in a cellular context.

## Results

### H2A.Z.1 slows the release of RNAPII towards progressive elongation

In vertebrates, H2A.Z is encoded by two independent genes *H2afz* (H2A.Z.1) and *H2afv* (H2A.Z.2 and its smaller spliceoform H2A.Z.2.2) that have distinct expression patterns and developmental roles^23,24,28^. Because H2A.Z.1 is strongly enriched at most promoters and is required for embryonic development as well as differentiation^17,19,22,29^, we focused on dissecting how this isoform regulates transcription in mouse embryonic stem cells (ESCs). We generated a homozygous ESC line containing a modified FKBP-V degron tag (dTAG) and 2x-HA epitope at the C-terminus of endogenous H2A.Z.1 (H2A.Z.1^dTAG^). This system allows for rapid and specific degradation of proteins by addition of the small molecule dTAG-13 (**Fig. 1a and Extended Data Fig. 1a,b**)^11^. Adding dTAG-13 to ESCs led to near complete loss of H2A.Z.1^dTAG^ within 8 hours as shown by immunoblot (**Fig. 1b and Extended Data Fig. 1c**). This system circumvents prior limitations using siRNA knock-down including the extensive time required to assess the consequences of depletion and possible off target effects. ChIP-seq with an anti-HA antibody revealed the highest distribution of H2A.Z.1^dTAG^ over promoters and some distal enhancers, correlating strongly with prior data both at a genome-wide (Spearman R = 0.89) and single gene level (**Extended Data Fig. 1d-f)**^19^. Moreover, H2A.Z.1^dTAG^ ESCs were able to form EBs (data not shown) indicating that the dTAG cassette does not interfere with variant incorporation or function. As expected, treating ESCs with dTAG-13 led to rapid and near complete loss of H2A.Z.1 ChIP-seq peaks across the genome within hours (**Fig. 1c and Extended Data Fig.1g**) validating the robustness of our system.

**Figure 1.**
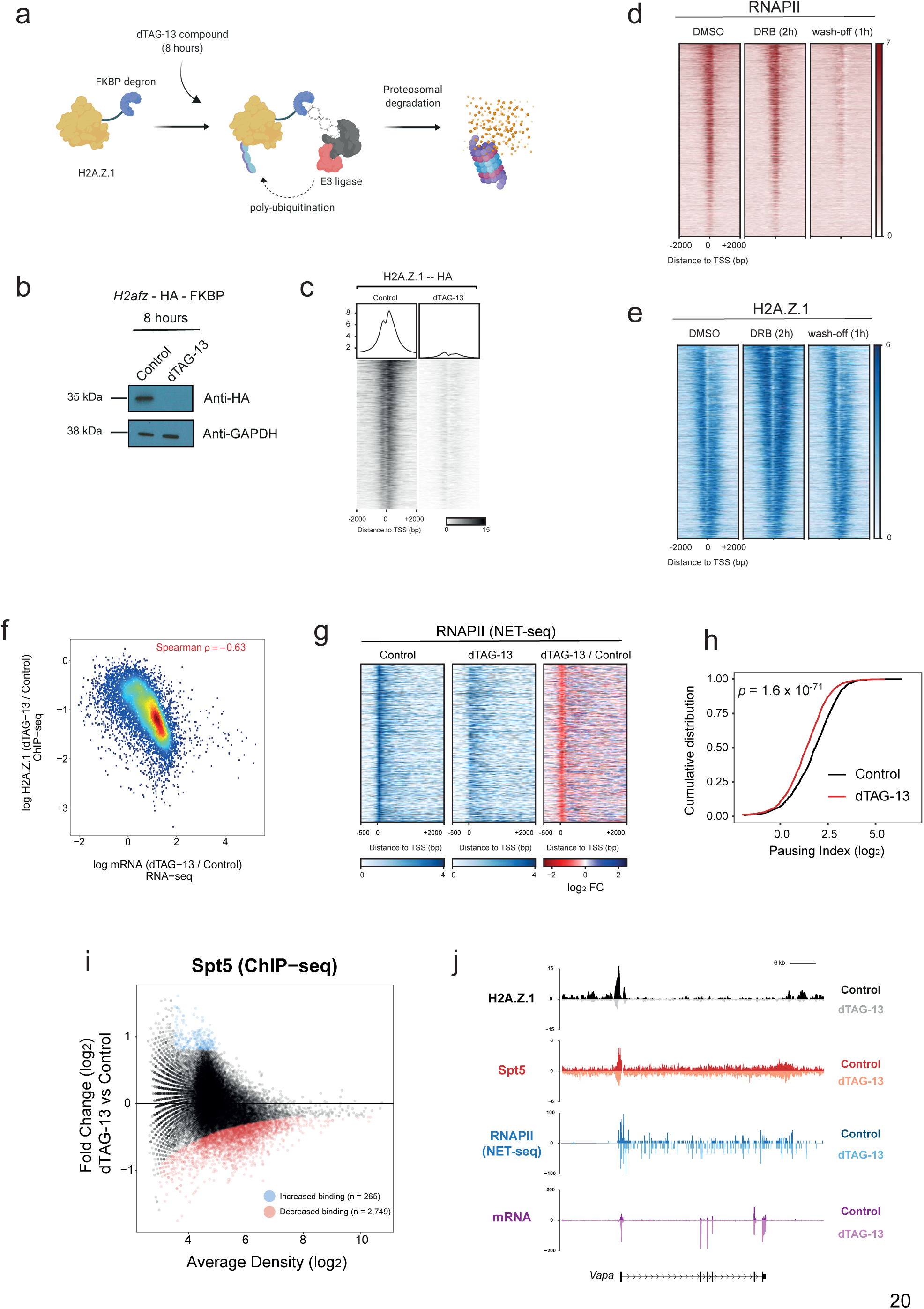
H2A.Z.1 attenuates the release of RNAPII towards progressive elongation. **a**. Schematic diagram of dTAG system for rapid depletion of H2A.Z.1 protein levels. **b**. Immunoblotting with anti-HA and anti-GAPDH antibodies in Control and dTAG-13 treated mESCs. **c**. ChIP-seq heatmaps over 7,789 protein-coding genes for H2A.Z.1 in control and dTAG-13 treated cells. **d**. ChIP-seq heatmaps over 7,624 uniquely annotated genes for RNAPII for Control (DMSO), DRB, and wash-off conditions. Genes are sorted by H3K4me3 levels. **e**. Same as (D) but for H2A.Z.1. **f**. Scatterplot of H2A.Z.1 ChIP-seq logFC (dTAG-13 / Control) versus mRNA (RNA-seq) logFC (dTAG-13 / Control). Spearman correlation in indicated in red (ρ = −0.63). **g**. Heatmap of RNAPII (NET-seq) for Control and H2A.Z.1-depleted (dTAG-13) mESCs over 4,184 uniquely annotated protein-coding genes. Genes are sorted by gene length. **h**. Cumulative distribution plot of RNAPII (NET-seq) Pausing Index for Control and H2A.Z.1-depleted (dTAG-13) mESCs for 4,184 uniquely annotated protein-coding genes. Significance was calculated using a paired Wilcoxon rank test. **i**. MA plot of Spt5 ChIP-seq between Control and dTAG-13 treated mESCs. Significant differential Spt5 binding displays either logFC < −0.2 or logFC > 0.2 and a P Value < 0.05. **j**. Single gene plots of ChIP-seq (H2A.Z.1 and Spt5), NET-seq (RNAPII), and RNA-seq over the *Vapa* gene. Control and dTAG-13 datasets are displayed in the positive and negative strands, respectively.

H2A.Z.1 is enriched at the promoters of most genes including both active (H3K4me3 only) and silent, bivalent (H3K4me3 and H3K27me3) genes in ESCs^17–19^. We observed lower H2A.Z.1 levels at genes with the highest RNAPII density by ChIP-seq (**Extended Data Fig. 2a**), consistent with more rapid nucleosome turnover during active transcription. To test this idea further, ESCs were treated with the rapid elongation inhibitor DRB (5,6-dichloro-1-beta-D-ribofuranosylbenzimidazole)^30^, an adenosine analogue that halts mRNA synthesis, followed by RNAPII and H2A.Z.1 ChIP-seq. DRB treatment for 2 hrs led to a decrease in the elongating form of RNAPII S2ph and increase in both RNAPII and H2A.Z.1 density at the promoter-proximal regions of active, but not bivalent genes (**Fig. 1d,e and Extended Data Fig. 2b-f**). Upon DRB wash off, H2A.Z.1 levels decreased at active promoters and RNAPII showed higher levels in gene bodies (**Fig. 1d,e and Extended Data Fig. 2b-d**). These data predict that dTAG-13 treatment will lead to increased transcript levels at genes with the highest H2A.Z.1 enrichment.

**Figure 2.**
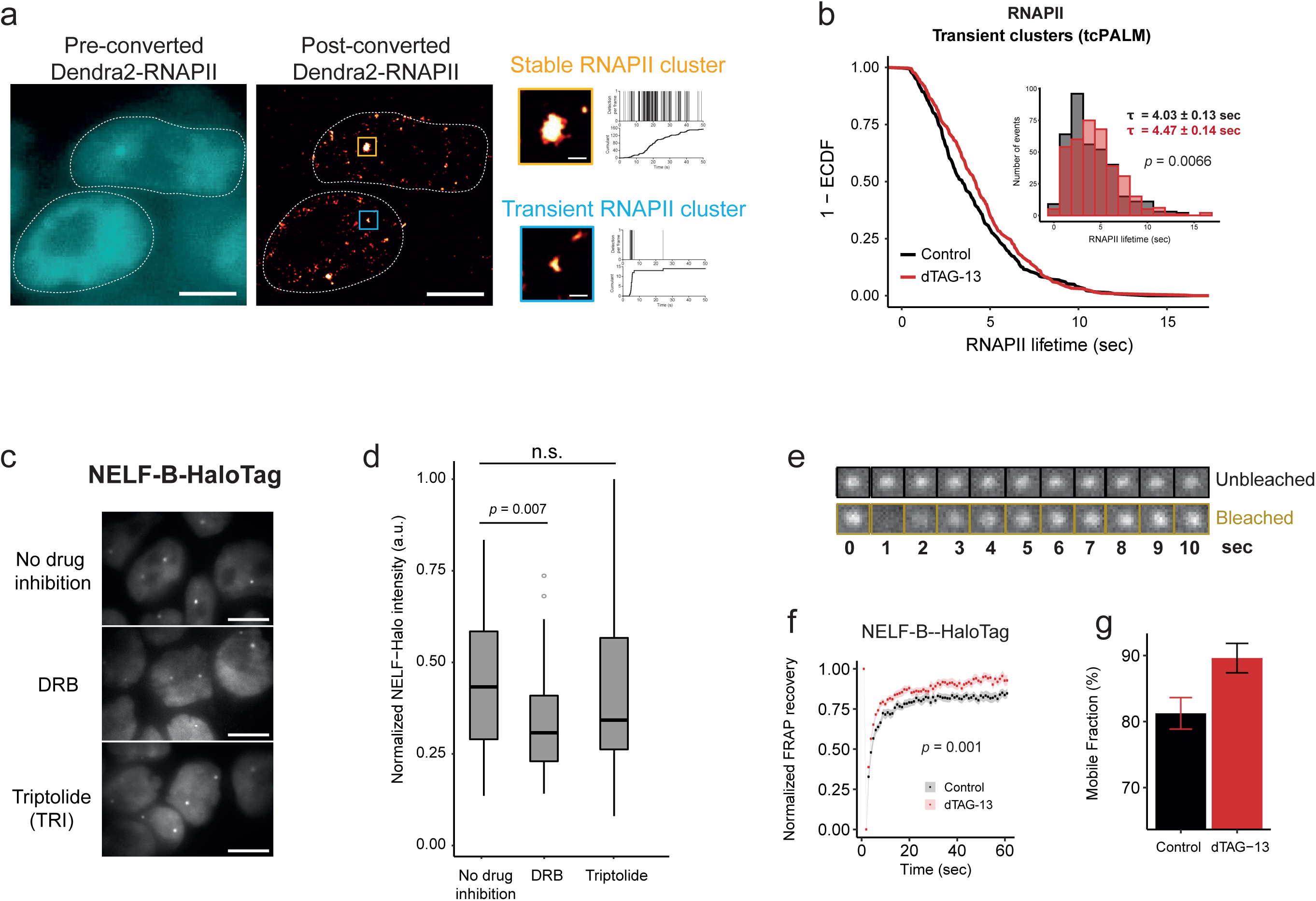
H2A.Z.1 controls NELF and RNAPII dynamics at a single molecule level. **a**. Live-cell direct image of pre-converted Dendra2-RNAPII (cyan) and super-resolution image of post-converted Dendra2-RNAPII (red) and corresponding time-correlated photoactivation localization microscopy (tcPALM) traces. Scale bar 5 µm and 5 nm (zoomed in clusters). **b**. Cumulative distribution and histogram of RNAPII transient cluster lifetime between Control (n = 360 clusters) and dTAG-13 (n = 363 clusters) measured by time correlated PALM (tcPALM). Significance is calculated using an unpaired Wilcoxon rank test. **c**. Live-cell direct images of NELF-Halo in Control and dTAG-13 conditions. Cells were treated with DMSO (no drug inhibition), DRB (50 µM), or Triptolide (50 µM) for 45 min. Scale bar 5 µm. **d**. Normalized intensity of NELF-Halo clusters displayed in **(c)**. No drug inhibition (n=42), DRB (n=35), TRI (n=56). Significance was calculated using an unpaired Wilcoxon rank test. **e**. FRAP analysis of NELF clusters. Images of a Halo-NELF cell before (0 sec), and immediately after (1-10 sec) bleaching. The black box indicates an unbleached control locus. The golden box indicates the position of the cluster on which the FRAP beam was focused. **f**. The normalized recovery curve for NELF (*n* = 23 cells) yielded a recovery fraction of ∼80% during the 60 sec observation in Control (black) conditions that increased to ∼90% in dTAG-13 treated cells. Dots and shaded areas represent mean and SEM values, respectively. The Welch unpaired t-test was used for significance. **g**. Mobile fraction of in Control (black) and dTAG-13 (red) conditions. Error bars represent ±SEM.

Using RNA-seq, indeed, we found that the vast majority of differentially expressed genes showed increased expression upon rapid H2A.Z.1 loss (**Extended Data Fig. 3a**). We observed a striking inverse correlation between H2A.Z.1 loss at promoters and increased expression of the corresponding gene (Spearman R=-0.63) (**Fig. 1f**). Coupled with our analysis showing an inverse correlation between H2A.Z.1 and RNAPII at promoters, these data suggest that H2A.Z.1 regulates RNAPII dynamics. To test this idea, we next performed Native Elongation Transcript Sequencing (NET-seq) to map the strand-specific location of RNAPII at high resolution^2^ in ESCs. Analysis of biological duplicates showed that RNAPII density was highly reproducible (Pearson’s R = 0.99 for both control and dTAG data) **(Extended Data Fig. 3b)**. Using available START-seq data in ESCs^3^, we compiled a list of 4,184 uniquely annotated transcription start sites (TSSs) corresponding to non-tandemly oriented protein-coding genes with detectable RNAPII levels (**Extended Data Fig. 3c**). Upon dTAG-13 treatment, RNAPII density significantly decreased at promoters and showed a concomitant increase in the gene body compared to controls (**Fig. 1g and Extended Data Fig. 3d**). We then calculated the RNAPII pausing index (PI) which is defined by the density of RNAPII over the promoter proximal region relative to the gene body^31^. A significant decrease (p = 1.6 x10^−71^) in the PI was observed upon H2A.Z.1 loss (**Fig. 1h**). ChIP-seq of Spt5, a subunit of the DSIF complex involved in promoter-proximal RNAPII pausing, also showed a global and gene-specific decrease upon dTAG-13 treatment (**Fig. 1i,j**). Together, these data suggest H2A.Z.1 acts as a barrier to transcription by hindering RNAPII pause-release.

**Figure 3.**
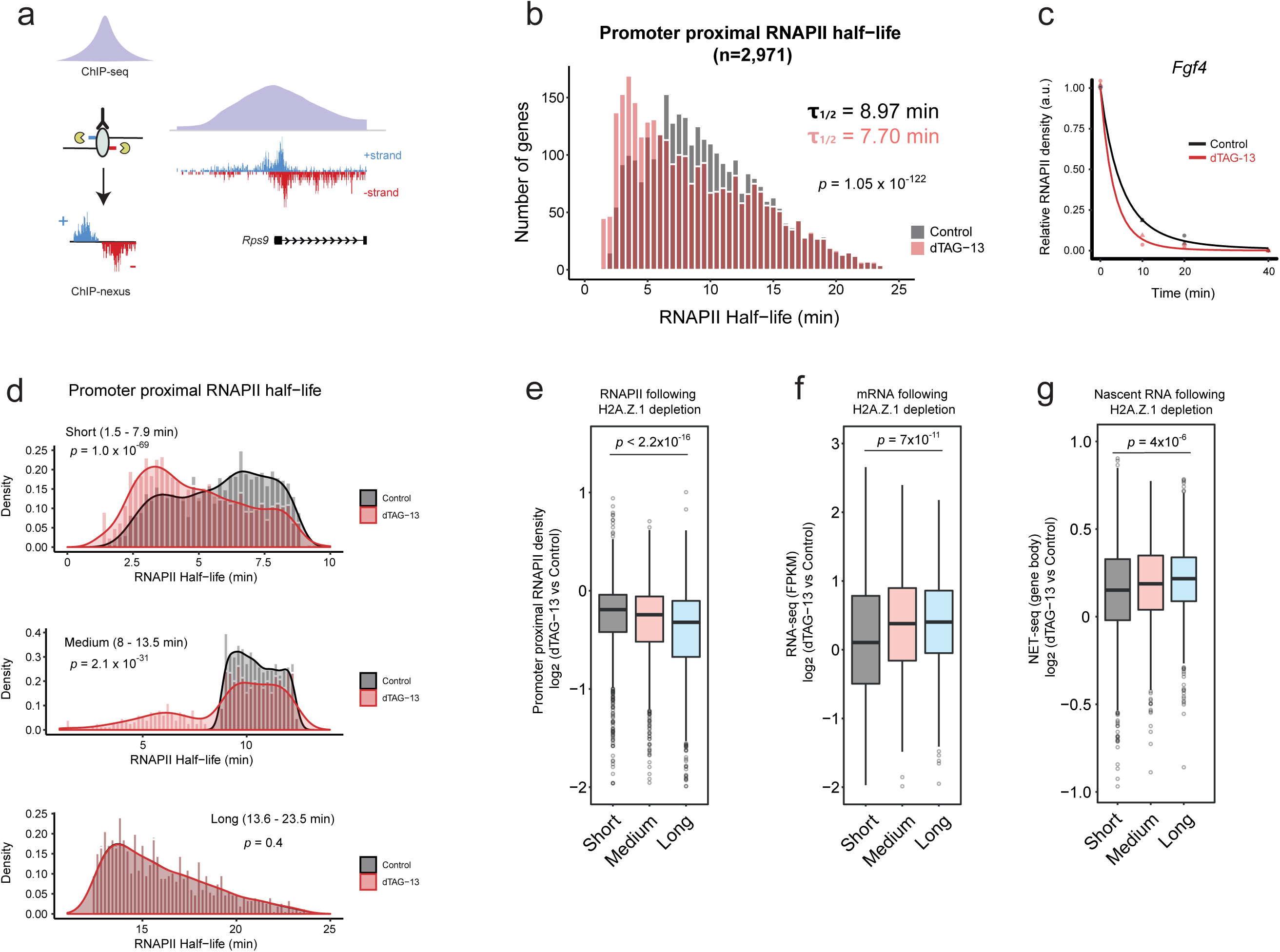
H2A.Z.1 controls RNAPII half-life of less stable promoters. **a**. Schematic diagram of the ChIP-nexus approach. Lambda (λ) exonuclease activity reveals transcription factor footprints at near nucleotide resolution. Representative tracks of RNAPII ChIP-seq and RNAPII ChIP-nexus over a single gene (*Rps9*). **b**. Histogram of the half-lives of paused RNAPII of 2,971 genes between Control (black) and dTAG-13 (red) conditions. Significance was calculated using a paired Wilcoxon rank test. **c**. The half-life of paused RNAPII was calculated on the basis of an exponential decay model. Normalized promoter proximal RNAPII density for the *Fgf4* gene is shown over the course of TRI treatment (0, 10, 20, and 40 min) both for Control (black) and dTAG-13 (red) conditions. **d**. Histograms of short (1.5-7.9 min), medium (8 −13.5 min), and long (13.6 - 23.5 min) RNAPII half-lives for Control and dTAG-13 conditions. Significance was calculated using a paired Wilcoxon rank test. **e**. Boxplots measuring log_2_ FC (dTAG-13 / Control) of promoter proximal RNAPII (ChIP-nexus) for the three different gene classes. Significance was calculated using an unpaired Wilcoxon rank test. **f**. Same as (E) but for mRNA levels (RNA-seq). **g**. Same as (E) but for gene body nascent RNA levels (NET-seq).

### H2A.Z.1 controls RNAPII and NELF dynamics at single molecule resolution

The above genomic approaches capture a snapshot of RNAPII at either a paused or elongating state, so we next wanted to quantify RNAPII dynamics at single molecule resolution in live cells. We engineered a photoconvertible Dendra2 tag on the largest subunit of endogenous RNAPII (RBP1) in H2A.Z.1^dTAG^ ESCs. ChIP-seq using an anti-Dendra2 antibody confirmed highest RNAPII enrichment at the TSSs of all protein-coding genes, both at a genome-wide and single gene level, strongly correlating with our NET-seq data (Pearson R = 0.75) (**Extended Data Fig. 4a-c**). Using single molecule super-resolution microscopy (tcPALM)^10,32^ in live H2A.Z.1^dTAG^;RNAPII^Dendra^ ESCs, we found that RNAPII exists both in stable and temporal clusters (∼100nm resolution) with an average cluster lifetime of 4.03 ± 0.13 sec (mean ± SEM of 360 clusters) for the latter clusters (**Fig. 2a**). Remarkably, H2A.Z.1 loss increased the average lifetime of temporal RNAPII clusters to 4.47 ± 0.14 sec (p=0.0066) (**Fig. 2b**). RNAPII cluster lifetime positively correlates with the rate of RNAPII re-initiation and ultimately mRNA output based on single molecule FISH^32–34^. In support of this correlation, treatment with the transcription initiation inhibitor triptolide (TRI)^35^ completely abolished RNAPII cluster dynamics in ESCs, whereas DRB treatment showed a less dramatic effect (**Extended Data Fig. 4d**).

**Figure 4.**
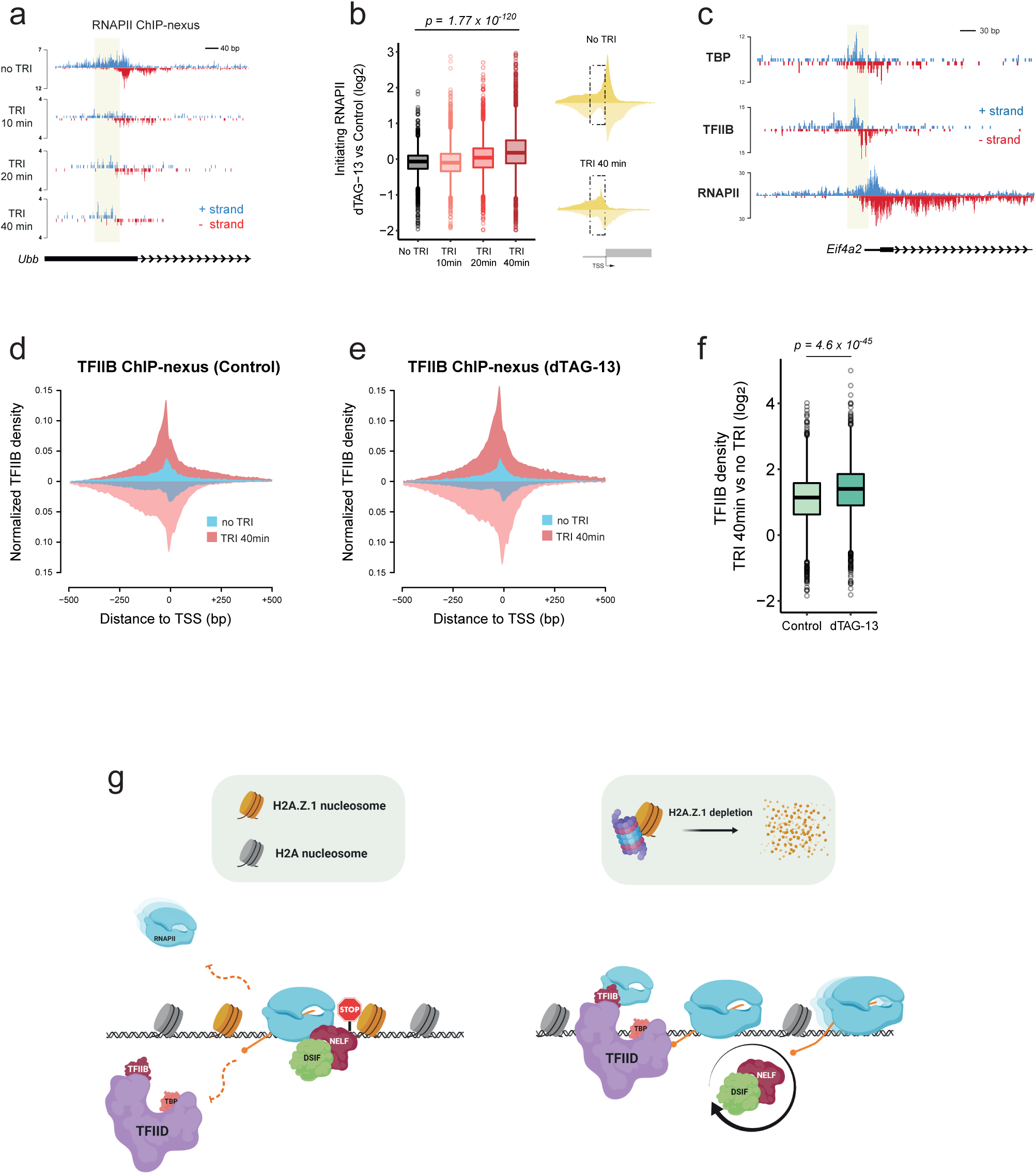
H2A.Z.1 controls Pre-initiation complex (PIC) recruitment at promoters. **a**. Single gene plot (*Ubb*) of RNAPII ChIP-nexus profiles over the course of Triptolide treatment (No TRI, 10 min, 20 min, and 40 min). RNAPII density is displayed both in the positive (blue) and negative (red) strand. Initiating RNAPII is highlighted in yellow. **b**. Boxplots measuring log_2_ FC (dTAG-13 vs Control) of initiating RNAPII over the course of TRI treatment. Significance was calculated using a Wilcoxon rank test. Schematic diagram displays ChIP-nexus RNAPII metaplots over the Transcription Start Site (TSS) whereas the dotted box denotes the area we used to quantify initiating RNAPII density (n = 3,298 genes). **c**. Single gene plot (*Eif4a2*) of TBP, TFIID, RNAPII ChIP-nexus profiles in Control conditions. ChIP-nexus density is displayed both in the positive (blue) and negative (red) strand. Initiating RNAPII, and PIC are highlighted in yellow. **d**. Metaplots of TFIIB ChIP-nexus in Control conditions with (“TRI 40min”) or without (“No TRI”) triptolide. **e**. Metaplots of TFIIB ChIP-nexus in H2A.Z.1-depleted (dTAG-13) conditions with (“TRI 40min”) or without (“No TRI”) triptolide. **f**. Boxplots measuring log_2_ FC (TRI 40min vs No TRI) TFIIB density between Control and dTAG-13 in an area 250 bp upstream of the TSS (n = 3,095 genes). Significance was calculated using a Wilcoxon rank test. **g**. Schematic diagram of how H2A.Z.1 controls proper execution of transcription programs. H2A.Z.1 blocks recruitment of the PIC and also widens the barrier for elongating RNAPII by promoting a more paused state (left panel). In the absence of H2A.Z.1, PIC is recruited faster while RNAPII is released in a faster rate from the promoter which is accompanied by faster NELF dynamics (right panel).

The significant decrease in the RNAPII pause index and Spt5/DSIF promoter levels upon dTAG-13 treatment as well as single molecule RNAPII cluster dynamics point to a direct role of H2A.Z.1 in regulating pause-release. Thus, we next measured the dynamics of other key pausing factors. Negative Elongation Factor (NELF) is a well characterized pausing complex that interacts dynamically with RNAPII upon promoter-release^4,16,36^. Thus, we analyzed available NELF ChIP-seq data in ESCs^31^ and found a strong overlap with H2A.Z.1 and RNAPII at active gene promoters (**Extended Data Fig. 4e**). To test the prediction that H2A.Z.1 effects NELF dynamics, we engineered a Halo-tag at the C-terminus of endogenous NELF-B, a subunit of the NELF complex, in H2A.Z.1^dTAG^ ESCs (**Extended Data Fig. 4f**). Live cell imaging of NELF-B-Halo showed bright punctate spots in the nucleus as previously described^37^ (**Fig. 2c**). To next measure NELF cluster dynamics relative to transcription, we treated ESCs for 45 min with either DRB or TRI to inhibit transcription elongation and initiation, respectively. NELF-B-Halo cluster size and intensity were unchanged by blocking initiation (**Fig. 2d**), consistent with the finding that NELF can be recruited to promoter-proximal regions through TFIID interactions^38^. In contrast, DRB treatment led to a significant decrease in NELF-Halo intensity (p<0.007) and dynamics by blocking elongation and subsequent re-loading or re-initiation of new RNAPII molecules^39^ (**Fig. 1d,e and Fig. 2d**) Thus, NELF clusters appear to mark regions of active transcriptional elongation in live cells.

These observations led us to test the dynamic relationship between H2A.Z.1 and NELF within these clusters. We performed fluorescence recovery after photobleaching (FRAP) in H2A.Z.1^dTAG^;NELF-B^Halo^ ESCs. FRAP showed that 81% of NELF was exchanged within ∼10 seconds in control ESCs similar to the very rapid turnover observed for RNAPII cluster dynamics (**Fig. 2e,f**)^10^. Remarkably, NELF clusters displayed a significantly higher recovery (89%; p=0.001) following dTAG-13 treatment suggesting NELF is more rapidly exchanged at active regions upon H2A.Z.1 loss (**Fig. 2f,g**). A similar trend was observed when performing FRAP in the presence of TRI (**Extended Data Fig. 4g,h**), consistent with our imaging data showing that blocking transcription initiation for 45 min does not decrease NELF cluster size due to elongation of engaged RNAPII. Together, measuring RNAPII and NELF dynamics in live cells suggest that H2A.Z.1 impacts both promoter-proximal pause-release and subsequent re-loading of RNAPII.

### H2A.Z.1 regulates promoter proximal RNAPII half-life

To test this idea that H2A.Z.1 prevents re-initiation presumably due to slowing the release of paused RNAPII at the gene level, we treated H2A.Z.1^dTAG^;RNAPII^Dendra2^ ESCs with TRI at different timepoints (10, 20, and 40 min) and performed ChIP-nexus^9^, a protocol that captures both initiating and stalled RNAPII at nucleotide resolution (**Fig. 3a**). As TRI treatment blocks new initiation^35^, paused RNAPII is eventually lost either by release of the elongation complex into the gene body or by premature termination of nascent transcripts^40^. As expected, TRI treatment caused a dramatic decrease of promoter-proximal RNAPII at active genes in both control and dTAG-13 treated ESCs (**Extended Data Fig. 5a,b**). ChIP-nexus also revealed a lower RNAPII pausing index upon H2A.Z.1 depletion compared to controls consistent with our NET-seq data (**Fig. 1h**) and with a more rapid RNAPII release into gene bodies (**Extended Data Fig. 5c**).

RNAPII half-life can be used as a measure of its turnover at gene promoters. Thus, we next fitted an exponential decay model using the time points of TRI treatment considering only those genes that maintain measurable RNAPII levels throughout the time course (see Methods). We calculated a median RNAPII half-life of 8.97 min in control ESCs (n = 2,971 genes), in good agreement with data in both *Drosophil*a^39,41^ and mammalian cell lines^40^. In contrast, RNAPII half-life decreased to 7.70 min (p = 1×10^−122^) upon dTAG-13 treatment (**Fig. 3b,c**). We then performed k-means clustering and divided genes into three categories based on RNAPII half-life/turnover in control ESCs: Short; 1.5-7.9 min (n = 1,449), Medium; 8-13.5 min (n = 730), and Long; 13.6 – 23.5 min (n = 728) (**Fig. 3d**). Following dTAG-13 treatment, RNAPII half-life was most dramatically reduced at genes displaying shorter half-lives (Short p = 1×10^−69^, Medium p = 2.1×10^−31^) (**Fig. 3d**). Surprisingly, although RNAPII half-life of more stably paused genes did not change appreciably upon H2A.Z.1 loss (Long p = 0.4), these genes exhibited a dramatic decrease of promoter-proximal RNAPII followed by a concomitant increase in nascent transcripts (**Fig. 3e-g**). Upon DRB treatment, we observed that genes with a longer RNAPII half-life also displayed a dramatic shift of RNAPII into the gene body upon release from the elongation block (**Extended Data Fig. 5d**). Moreover, H2A.Z.1 slows pause-release and nascent transcription at stably paused genes, a class of genes that appear most sensitive to signalling cues^3,39^. Notably, these stably paused genes show lower expression in ESCs and have higher promoter GC content (**Extended Data Fig. 5d-g**), a sequence feature associated with stable RNAPII pausing in metazoans^42,43^. Thus, limiting pause release and subsequent RNAPII loading may safeguard proper activation of H2A.Z.1 genes.

### H2A.Z.1 controls Pre-initiation complex (PIC) recruitment at promoters

Our observations suggest that stably paused genes experience more rapid reloading of RNAPII upon H2A.Z.1 depletion. Although our tcPALM data are consistent with an overall increase in initiating RNAPII upon dTAG-13 treatment (**Fig. 2b**), we wanted to measure RNAPII PIC recruitment and initiation at individual promoters. RNAPII ChIP-nexus in combination with TRI treatment showed a dramatic increase in initiating Serine 5 phosphorylated CTD levels (RNAPII S5ph) (**Extended Data Fig. 6a**) and shift of RNAPII occupancy upstream of the TSS (**Fig. 4a**) consistent with the position of the PIC^39,44^. Notably, after 40 min of TRI treatment, we observed similarly higher initiating RNAPII levels in dTAG-13 treated compared to control ESCs (**Fig. 4b and Extended Data Fig. 6b,c**).

To validate that the increase in initiating RNAPII upon H2A.Z.1 loss is due to new PIC recruitment, we next performed ChIP-nexus for the PIC subunits TFIIB and TBP to map their location at base pair resolution. Both TFIIB and TBP displayed a strong enrichment ∼20-30bp upstream of the TSS with high reproducibility between biological replicates (**Fig. 4c and Extended Data Fig. 7a,b**). *In vitro*, PIC dynamics is on the order of seconds^14,45–47^ and promoter proximal paused RNAPII is refractory to new transcription initiation^39^. To first measure new TFIIB and TBP enrichment, we treated cells with TRI for 40 min to enable complete release of paused RNAPII. ChIP-nexus revealed an increase of new PIC recruitment at RNAPII-dependent promoters either upon TRI and/or dTAG-13 treatment (**Fig. 4d-f and Extended Data Fig. 7c-e**) with the most dramatic increase observed at stably paused (Long) promoters (**Extended Data Fig. 7f**). Collectively, our data support a model whereby H2A.Z.1 enables multistep control of transcription by slowing RNAPII pause-release which subsequently inhibits PIC assembly and re-initiation.

RNAPII pausing is critical to allow genes to appropriately respond to developmental and environmental signals by controlling the timing, rate, and magnitude of the transcriptional response^3,44,48^. In addition to known pausing factors, our quantitative molecular analysis provides strong evidence that H2A.Z.1 acts as a barrier to transcription, affecting both RNAPII pause-release and re-engagement of new RNAPII at promoters in mammalian cells (**Fig. 4g**). Thus, H2A.Z.1 likely coordinates synchronous gene expression in response to signaling cues. Consistent with this idea, stably paused genes show more synchronous expression, lower cell to cell variability, and are typically signal-responsive genes^49^. Thus, H2A.Z.1 may serve as a GO or NO-GO decision point whereby RNAPII can either undergo early transcript termination or progressive elongation depending on cellular signals. Notably, in the absence of NELF, RNAPII experiences a second pause around the +1 nucleosome dyad^16^. Our data argue that site-specific H2A.Z incorporation surrounding the NDR is critical for this additional pause. Thus, our data also predict that H2A.Z.1 nucleosomes upstream of the TSS play a role in regulating PIC assembly and re-initiation. Together, our work uncovers a mechanistic link between H2A.Z.1 and transcription dynamics and provides an explanation for its promoter enrichment at RNAPII-regulated genes and requirement in metazoan development.

## Acknowledgements

We thank the Boyer lab, Seychelle Vos, Eliezer Calo and Craig Peterson for helpful discussions and insightful comments on the manuscript. This work was supported by NIGMS R01-GM134734 to I.C., NHLBI R01-HL140471 to L.A.B. and the Koch Institute Core Grant P30-CA14051 from the National Cancer Institute.

## Accession Numbers

Raw and normalized sequencing data has been deposited under GEO accession number GSEXXXXXXX.

## Author Contributions

CM and LAB designed the study. CM performed experiments and analyzed data. AA assisted with FRAP image acquisition and analysis. CL and IC assisted with tcPALM. CM and LAB wrote the manuscript with input from all authors.

## Materials and Methods

### ESC culture

V6.5 murine embryonic stem cells (mESC) were cultured in 2i + LIF conditions^50^ for all assays. Genome editing was performed in serum + LIF conditions. Cells were cultured at 37°C, 5% CO2, on 0.1% gelatin coated tissue culture plates. The media used for general culturing in serum + LIF conditions is as follows: DMEM-KO (Invitrogen 10829-018) supplemented with 15% fetal bovine serum (Hyclone characterized SH3007103), 1,000 U/ml LIF, 100 mM nonessential amino acids (Invitrogen 11140-050), 2 mM L-glutamine (Invitrogen 25030-081), 100 U/mL penicillin, 100 mg/mL streptomycin (Invitrogen 15140-122), and 8 ul/mL of 2-mercaptoethanol (Sigma M7522). The media used for 2i + LIF media conditions is as follows: 484 mL DMEM/F12 (GIBCO 11320), 2.5 mL N2 supplement (GIBCO 17502048), 5 mL B27 supplement (GIBCO 17504044), 0.5 mM L-glutamine (GIBCO 25030), 1X non-essential amino acids (GIBCO 11140), 100 U/mL Penicillin-Streptomycin (GIBCO 15140), 0.1 mM β-mercaptoethanol (Sigma M6250), 1 uM PD0325901 (Stemgent 04-0006), 3 uM CHIR99021 (Stemgent 04-0004), and 1000 U/mL recombinant LIF.

### Immunoblots

To assay immunoprecipitation results by western blot, 500ng of each sample was run on a 4%– 20% Bis-Tris gel (Bio-rad 3450124) using SDS running buffer (Bio-rad 1610788) at 120V for 10 minutes, followed by 180V until dye front reached the end of the gel. Protein was then wet transferred to a nitrocellulose membrane using the Trans-blot turbo transfer system (Bio-rad). After transfer, the membrane was blocked with 5% BSA for 1h on a rotor.

Membrane was then incubated with 1:5000 Anti-HA (Cell Signaling 3724), 1:1000 Anti-Nelf-B (Cell Signaling 14894), 1:1000 Anti-RNAPII S5ph (Active Motif 39750), 1:1000 Anti-RNAPII S2ph (Active Motif 39564), 1:1000 Anti-GAPDH (Cell Signaling 8884) diluted in 5% BSA in TBST and incubated for 2h at room temperature. Following several washes with TBST, membranes were incubated either with 1:10,000 Anti-mouse HRP (Cell Signaling 7076) or with 1:10,000 Anti-rabbit HRP (R&D Systems FIN1818061) for 1h at room temperature with rotation. After extensive wash with TBST, membranes were developed with ECL substrate (Bio-rad 1705060) and imaged using Azure 600 or exposed using film.

### Chromatin Immunoprecipitation (ChIP)

ChIP was performed using ∼10 million ESCs per assay as previously described^51^. Briefly, cells were cross-linked with 1% formaldehyde for 10 min followed by 5 min quenching with 125 mM glycine. After washing with PBS buffer, the cells were collected and lysed in cell lysis buffer (5 mM Tris, pH 8.0, 85 mM KCl, and 0.5% NP-40) with x1 Halt Protease Inhibitor cocktail (ThermoFisher 87786) and 1mM PMSF (Sigma 10837091001). Pellets were spun for 5 min at 6000 rpm at 4°C. Nuclei were lysed in nuclei lysis buffer (1% SDS, 10 mM EDTA, and 50 mM Tris–HCl) and samples were sonicated for 12 min on a Covaris Sonicator. The samples were centrifuged for 20 min at 13,000 rpm at 4°C and the supernatant was diluted in IP buffer (0.01% SDS, 1.1% Triton-X-100, 1.2 mM EDTA, 16.7 mM Tris–HCl, and 167 mM NaCl), and the appropriate antibody (10 μg) was added and incubated overnight at 4°C with rotation. Antibodies used in this study; Anti-HA (Cell Signaling 3724), Anti-Spt5 (Santa Cruz sc-133217), Anti-Dendra2 (OriGene TA180094), Anti-TFIIB (Cell Signaling 4169), Anti-TBP (Abcam, ab28175). Two biological replicates were prepared for each condition using independent cell cultures and chromatin precipitations. Following overnight incubation, 50 μl Protein G Dynabeads (Life Technologies 10009D) were added for 1 h at room temperature with rotation. Beads were washed once for 1 min with rotation with each of the following buffers: Low salt buffer (0.1% SDS, 1% Triton-X-100, 2mM EDTA, 20mM Tris pH 8.0, 150mM NaCl), High salt buffer (0.1% SDS, 1% Triton-X-100, 2mM EDTA, 20mM Tris pH 8.0, 500mM NaCl), LiCl buffer (0.25M LiCL, 1% NP-40, 1% dioxycholate, 10mM Tris pH 8.0, 1mM EDTA), and TE buffer (50mM Tris pH 8.0, 10mM EDTA). DNA was eluted off the beads by rotation at room temperature for 30 min in 200 μL elution buffer (1% SDS, 0.1M NaHCO3). Cross-links were reversed at 65°C for 4h. RNA was degraded by the addition of 3 μL of 33 mg/mL RNase A (Sigma R4642) and incubation at 37°C for 2 hours. Protein was degraded by the addition of 5 μL of 20 mg/mL proteinase K (Invitrogen 25530049) and incubation at 50°C for 1 hour. Phenol:chloroform:isoamyl alcohol extraction was performed followed by ethanol precipitation, the resulting DNA was resuspended in 20 μL H2O, and used for either qPCR or sequencing.

For ChIP-seq experiments, purified 10-20 ng of ChIP DNA was used to prepare Illumina multiplexed sequencing libraries. Libraries for Illumina sequencing were prepared following the NEBNext DNA Library Prep Master Mix kit (NEB E6040). Amplified libraries were size selected using a 2% agarose gel to extract fragments between 200 and 600 bp.

### ChIP-nexus

For each ChIP we used ∼20 million crosslinked mouse ESCs spiked-in with 5% human U2OS cells expressing Dendra2-RNAPII to account for loss of RNAPII during triptolide treatment. For TBP and TFIIB ChIPs, mouse ESCs were spiked-in with 5% human iPSCs cells. ChIP was performed as described above. The ChIP-nexus libraries were constructed as previously described with minor modifications^9^. IPed-bead bound material was End-repaired, followed by dA-tailing and adaptor ligation. The ChIP-nexus adaptor mix contained four fixed barcodes (ACTG, CTGA, GACT, TGAC). Barcode was extended with Phi29 polymerase, followed by λ exonuclease digestion, ethanol precipitation and ssDNA circularization. A detailed ChIP-nexus protocol can be found online (<u>ChIP-nexus protocol v.2019</u>). At least two biological replicates were performed for each factor to obtain coverage of at least 80 million reads per condition. Single-end sequencing of 75 bp was performed on an Illumina NextSeq 500 instrument.

### ChIP-seq/nexus analysis

For ChIP-seq, the samples were single-end deep-sequenced and reads were aligned to the mm10 genome using Bowtie2 (v 2.3.5)^52^. Peak calling was performed using PePr (v 1.1)^53^ with peaks displaying an FDR < 10^−5^ considered statistically significant. All aligned ChIP-seq BAM files were converted to bigwig (10 bp bin) and normalized to 1× sequencing depth using deepTools (v 3.0)^54^. Blacklisted mm10 coordinates were further removed from the analysis. Average binding profiles were visualized using R (v 3.6.1). Heatmaps were generated with deepTools.

For ChIP-nexus, fastq files were filtered for the presence of the four mixed barcodes and were deduplicated using Q-Nexus^55^. Bowtie2 (v 2.3.5) was used for alignment to either the mm10 or hg19 genome. Normalization factors were computed for reads mapping uniquely to the human genome using Deseq2^56^. All aligned mouse ChIP-nexus BAM files were converted to bigwig (1 bp bin), separated by strand, and normalized to human spike-in controls using deepTools (v 3.0) with an “–offset 1” to record the position of the 5’ end of the sequencing read which corresponds to TF footprint.

To calculate the half-life of paused RNAPII for each gene, RNAPII density was calculated in a 250-bp window around the TSS. RNAPII time-course measurements were fitted into an exponential decay model using the “RNAdecay” ^57^ R package. We selected genes fulfilling the current criteria; 1) detectable RNAPII levels (RPM > 0.1), 2) highest RNAPII density at “No TRI” condition, and 3) low variance between replicates (sigma < 0.05). Genes fitting the above criteria (n = 2,971) were used to calculate RNAPII half-life. The full list of genes is displayed in **Extended Data Table 1**.

### Genome Editing using CRISPR/Cas9

The CRISPR/Cas9 system was used to genetically engineer ESC lines. Target-specific oligonucleotides were cloned into the pSpCas9-GFP plasmid (Addgene PX458) using BbsI restriction digestion. The oligo sequences used for cloning are;

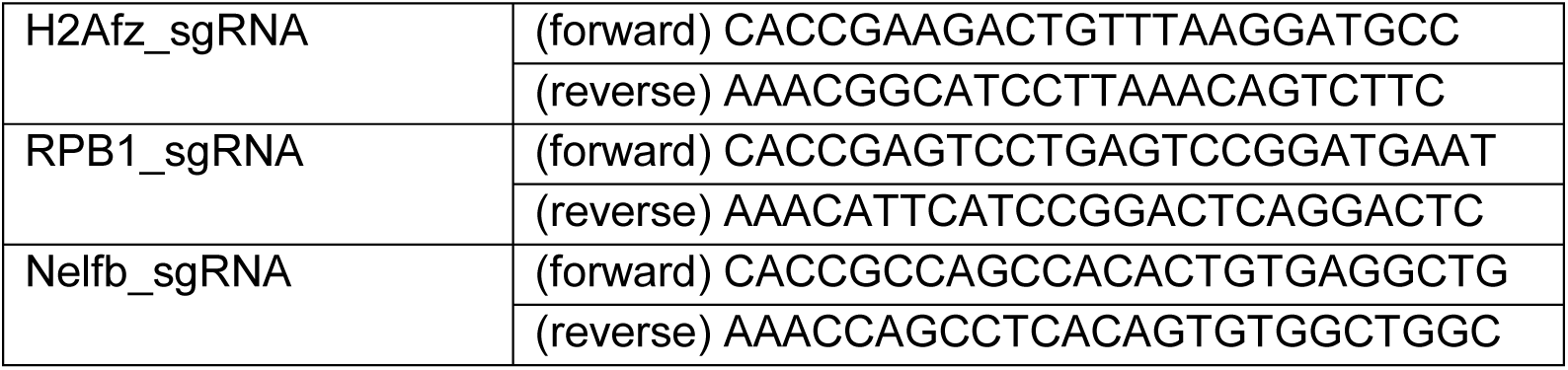

Plasmids containing H2afz-FKBP.knock-in.BFP and Nelfb-Halo DNA repair templates were synthesized with long homology arms (∼800 bp) by Genewiz FragmentGene and assembled using the NEBuilder HiFi DNA Assembly Cloning kit (NEB E5520S). The Dendra2-RBP1 plasmid and guide were used as previously described^10^.

For the generation of the endogenously tagged lines, 1 million mES cells were transfected with 1.25 μg Cas9 plasmid and 1.25 μg non-linearized repair template plasmid. Cells were sorted after 30 hours for the presence of Cas9-GFP. Cells were expanded for five days and then sorted again either for BFP (H2afz-FKBP) or GFP (RBP1-Dendra2). Cells that were transfected with the Nelfb-Halo repair template were incubated for 15 min with 500nM Halo TMR ligand (Promega G8251), washed 3 times with 2i media and further incubated for another 30 min before sorting for Texas Red. Sorted cells were plated at a low density (∼400 cells) on 10cm plates with irradiated MEFs and grown for approximately one week. Individual colonies were picked using a stereoscope into a 96-well plate. Cells were expanded and genotyped by PCR. Clones with a homozygous knock-in tag were further expanded and used for experiments.

### Super-resolution imaging

Live cell PALM imaging was carried out as described before^10^. Briefly, cells were incubated in L-15 medium and were simultaneously illuminated with 1.3 W/cm2 near UV light (405nm) for photoconversion of Dendra2 and 3.2 kW/cm2 (561nm) for fluorescence detection with an exposure time of 50ms. We acquired images of Dendra2-RNAPII for 50s (1000 frames) for quantification of transient clusters. Super-resolution images were reconstructed using MTT ^58^ and qSR^59^. For imaging NELFb-Halo clusters, cells were incubated for 15 min with 500nM Halo TMR ligand (Promega G8251), washed with 2i media three times followed by 30 min incubation in 2i media without the ligand to remove unbound HaloTaq ligands before fluorescence imaging in L-15 medium. We acquired 100 frames (5s) with 561nm excitation.

### Density based spatial clustering of applications with noise (DBSCAN) analysis

We used a clustering algorithm, density based spatial clustering of applications with noise (DBSCAN), to define the area of clustered regions in super-resolution data, as previously described^10^. Based on the single molecule localization in a super-resolution image, DBSCAN tests if a localization can be grouped with nearby localizations. Once a localization has a minimum number of nearby localizations (N) within a specific distance (R), it is grouped with other localizations also satisfying N and R criteria in their local neighbourhood. We defined the parameters N = 25-30 (points) and R = 90-95 (nm) for Dendra2-RNAPIII clustering. Varying the parameters for an image, we chose a set of parameters (N, R) for which the DBSCAN clustering result agrees with intensified regions in the super-resolution image generated using the qSR software module. For DBSCAN analysis, we used the ‘DBSCAN’ module embedded in the qSR software.

### Fluorescence Recovery After Photobleaching (FRAP)

FRAP experiments were carried out as previously described^10^. Images were taken at a single confocal plane with an exposure time of 200ms. The bleach spot was taken at the center of each cluster and images were acquired at 1s intervals for 1min. Imaging was done on an Andor Revolution Spinning Disk Confocal microscope was used with the FRAPPA module. To quantify FRAP recovery, photobleaching was corrected by normalizing to non-bleached areas using the FIJI plugin FRAP profiler (http://worms.zoology.wisc.edu/research/4d/4d).

### RNA-isolation and sequencing

RNA was isolated using the RNeasy Plus Mini Kit (QIAGEN 74136) according to manufacturer’s instructions. ERCC spike-in controls were added to the RNA samples prior to library preparation. Stranded Ribo-depleted selected libraries were prepared using the TruSeq Stranded mRNA Library Prep Kit (Illumina RS-122-2101) according to manufacturer’s standard protocol. Libraries were subject to 75 bp paired-end end sequencing on an Illumina NextSeq instrument.

### RNA-seq analysis

Sequenced reads were aligned to the mm10 genome via STAR (v 2.7.2b)^60^. Gene counts were obtained from featureCounts of the Rsubread package (R/Bioconductor). Only genes with CPM

> 2 were included in subsequent analysis. Normalization factors from ERCC spike-in controls were calculated using edgeR^61^ and applied to the counts mapping to the mm10 genome. Differential expression was performed using the limma^62^ package. Significant genes with an absolute Fold change > 1.5 and adjusted *P*-value < 0.05 were considered as differentially expressed (**Extended Data Table 2**).

### Cell fractionation, RNA preparation and sequencing library construction for NET-seq

The cell fractionation was performed as described previously^63^. 10 million ES cells are washed with 500 ml of pre-cooled 1x PBS, resuspended in 150 ml Cytoplasmic lysis buffer (0.15% (v/v) NP-40, 10 mM Tris-HCl (pH 7.0), 150 mM NaCl, 25 mM a-amanitin (MCE HY-19610), 50 U SUPERase.In (Life Technologies AM2694), x1 Halt Protease Inhibitor cocktail mix (ThermoFisher 87786) and incubated on ice for 5 min. The cell lysate is layered over 400 ml of Sucrose buffer (10 mM Tris-HCl (pH 7.0), 150 mM NaCl, 25% (w/v) sucrose, 25 mM a-amanitin, 50 U SUPERase.In, x1 Halt Protease Inhibitor cocktail mix) and centrifuged at 16,000 g for 10 min at 4C. The nuclei pellet is resuspended in 500 ml Nuclei wash buffer (0.1% (v/v) Triton X-100, 1 mM EDTA, in 1x PBS, 25 mM a-amanitin, 50 U SUPERase.In, 1x Protease inhibitor mix) and centrifuged at 1,150 g for 1 min at 4C. Washed nuclei are resuspended in 200 ml Glycerol buffer (20 mM Tris-HCl (pH 8.0), 75 mM NaCl, 0.5 mM EDTA, 50% (v/v) glycerol, 0.85 mM DTT, 25 mM a-amanitin, 50 U SUPERase.In, 1x Protease inhibitor mix). Next, 200 ml of Nuclei lysis buffer (1% (v/v) NP-40, 20 mM HEPES pH 7.5, 300 mM NaCl, 1M Urea, 0.2 mM EDTA, 1 mM DTT, 25 mM a-amanitin, 50 U SUPERase.In, 1x Protease inhibitor mix) are added, mixed by pulsed vortexing and incubated on ice for 2 min. The lysate is centrifuged at 18,500 g for 2 min at 4C. The chromatin pellet is resuspended in 50 ml Chromatin resuspension solution (25 mM a-amanitin, 50 Units SUPERase.In, 1x Protease inhibitor mix in 1x PBS) before RNA preparation. Biological replicates were obtained from two independent Control and dTAG-13 treated cells. Prior to library preparation, SIRV spike-in controls (1:10,000 dilution) were added to the extracted RNA to account for loss of RNA during library preparation. NET-seq library construction was conducted as originally described^50^.

More than 90% recovery of ligated RNA and cDNA was achieved from 15% TBE-Urea (Invitrogen EC6885BOX) and 10% TBE-Urea (Invitrogen EC6875BOX), respectively, by adding RNA recovery buffer (R1070-1-10; Zymo Research) to the excised gel slices and further incubating at 70°C (1,500 rpm) for 15 min. Gel slurry was added to a ZymoSpin IV Column (Zymo Research C1007-50) and cDNA was precipitated for subsequent library preparation steps. cDNA containing the 3’ end sequences of a subset of mature and heavily sequenced snRNAs, snoRNAs, and rRNAs were specifically depleted using biotinylated DNA oligos^50^. Oligo-depleted circularized cDNA was amplified by PCR and double-stranded DNA was run on a 3% low melt agarose gel. The final NET-seq library running at ∼150 bp was extracted and further purified using the ZymoClean Gel DNA recovery kit (Zymo Research D4007). Sample purity and concentration was assessed in a 2200 TapeStation and further deep sequenced in a HiSeq 2500 Illumina Platform.

### Processing and alignment of NET-seq reads

All the NET-seq FASTQ files were processed using custom Python scripts (https://github.com/BradnerLab/netseq). The sequencing reads are aligned to the mouse reference genome (mm10) using the STAR (v 2.7.2b) aligner^60^. PCR duplicates and reads arising from RT bias were also removed. Reads mapping exactly to the last nucleotide of each intron and exon (splicing intermediates) were further removed from the analysis. The final NET-seq BAM files were converted to bigwig, separated by strand, and normalized to SIRV spike-in controls using deepTools (v 3.0) with an “–offset 1” to record the position of the 5’ end of the sequencing read which corresponds to the 3’ end of the nascent RNA. NET-seq tags sharing the same or opposite orientation with the TSS were assigned as “sense” and “antisense” tags, respectively. Promoter-proximal regions were carefully selected for analysis to ensure that there is minimal contamination from transcription arising from other transcription units. Genes overlapping within a region of 2 kb upstream of the TSS were removed from the analysis.

### RNAPII pausing index calculation

The RNAP II pausing index is determined by dividing the coverage in the region −30 to +250 bp around transcription start sites by the coverage in the region +300 bp downstream of the TSS to the transcription end site. The analysis was performed for non-overlapping protein-coding genes displaying a detectable signal of RNAPII (RPM > 0.1) at the promoter proximal region.

### Publicly available ChIP-seq data

H3K4me3;H3K27Ac;H3K27me3 (GSE47950)^64^, H2A.Z.1-GFP (GSE40063)^19^, RING1B (GSE69955)^65^. *Bona fide* active (H3K4me3 only) and bivalent (H3K4me3 & H3K27me3) gene co-ordinates were downloaded from Mas *et al* 2018^66^.

## References

1. Kwak, H., Fuda, N. J., Core, L. J. & Lis, J. T. Precise Maps of RNA Polymerase Reveal How Promoters Direct Initiation and Pausing. Science 339, 950–953 (2013).

2. Mayer, A. et al. Native elongating transcript sequencing reveals human transcriptional activity at nucleotide resolution. Cell 161, 541–544 (2015).

3. Williams, L. H. et al. Pausing of RNA Polymerase II Regulates Mammalian Developmental Potential through Control of Signaling Networks. Mol. Cell 58, 311–322 (2015).

4. Lee, C. et al. NELF and GAGA Factor Are Linked to Promoter-Proximal Pausing at Many Genes in Drosophila. Mol. Cell. Biol. 28, 3290–3300 (2008).

5. Jonkers, I. & Lis, J. T. Getting up to speed with transcription elongation by RNA polymerase II. Nat Rev Mol Cell Biol 16, 167–177 (2015).

6. Chen, Z. et al. High-resolution and high-accuracy topographic and transcriptional maps of the nucleosome barrier. Elife 8:e48281 (2019).

7. Weber, C. M., Ramachandran, S. & Henikoff, S. Nucleosomes are context-specific, H2A.Z-Modulated barriers to RNA polymerase. Mol. Cell 53, 819–830 (2014).

8. Jang, C. W., Shibata, Y., Starmer, J., Yee, D. & Magnuson, T. Histone H3.3 maintains genome integrity during mammalian development. Genes Dev. 29, 1377–1393 (2015).

9. He, Q., Johnston, J. & Zeitlinger, J. ChIP-nexus enables improved detection of in vivo transcription factor binding footprints. Nat. Biotechnol. 33, 395–401 (2015).

10. Cho, W.-K. et al. Mediator and RNA polymerase II clusters associate in transcription-dependent condensates. Science 6400, 412–415 (2018).

11. Nabet, B. et al. The dTAG system for immediate and target-specific protein degradation. Nat. Chem. Biol. 14, 431–441 (2018).

12. Core, L. & Adelman, K. Promoter-proximal pausing of RNA polymerase II: a nexus of gene regulation. Genes Dev. 33, 960–982 (2019).

13. Vos, S. M., Farnung, L., Urlaub, H. & Cramer, P. Structure of paused transcription complex Pol II–DSIF–NELF. Nature 560, 601–606 (2018).

14. Sainsbury, S., Bernecky, C. & Cramer, P. Structural basis of transcription initiation by RNA polymerase II. Nat Rev Mol Cell Biol 16, 129–143 (2015).

15. Yamaguchi, Y. et al. NELF, a Multisubunit Complex Containing RD, Cooperates with DSIF to Repress RNA Polymerase II Elongation. Cell 97, 41–51 (1999).

16. Aoi Y, Smith ER, Shah AP, et al. NELF Regulates a Promoter-Proximal Step Distinct from RNA Pol II Pause-Release. Mol Cell 78(2), 261–274 (2020).

17. Hu, G. et al. H2A.Z facilitates access of active and repressive complexes to chromatin in embryonic stem cell self-renewal and differentiation. Cell Stem Cell 12, 180–192 (2013).

18. Ku, M. et al. H2A.Z landscapes and dual modifications in pluripotent and multipotent stem cells underlie complex genome regulatory functions. Genome Biol. 13(10) R85 (2012).

19. Subramanian, V. et al. H2A.Z Acidic Patch Couples Chromatin Dynamics to Regulation of Gene Expression Programs during ESC Differentiation. PLoS Genet. 9 (8) (2013).

20. Maze, I., Noh, K.-M., Soshnev, A. A. & Allis, C. D. Every amino acid matters: essential contributions of histone variants to mammalian development and disease. Nat. Rev. Genet. 15, 259–271 (2014).

21. Subramanian, V., Fields, P. A. & Boyer, L. A. H2A.Z: a molecular rheostat for transcriptional control. F1000Prime Rep. 7 (01), (2015).

22. Faast, R. et al. Histone variant H2A.Z is required for early mammalian development. Curr. Biol. 11, 1183–1187 (2001).

23. Greenberg, R. S., Long, H. K., Swigut, T. & Wysocka, J. Single Amino Acid Change Underlies Distinct Roles of H2A.Z Subtypes in Human Syndrome. Cell 178, 1421–1436 (2019).

24. Lamaa, A. et al. Integrated analysis of H2A.Z isoforms function reveals a complex interplay in gene regulation. Elife 9, e53375 (2020).

25. Reschen, M. et al. Floating–Harbor syndrome and polycystic kidneys associated with SRCAP mutation. Am. J. Med. Genet. Part A 158A, 3196–3200 (2012).

26. Chen, I. Y. et al. Histone H2A.z is essential for cardiac myocyte hypertrophy but opposed by silent information regulator 2a. J. Biol. Chem. 281, 19369–19377 (2006).

27. Svotelis, A., Gevry, N., Grondin, G. & Gaudreau, L. H2A.Z overexpression promotes cellular proliferation of breast cancer cells. Cell Cycle 9, 364–370 (2010).

28. Dryhurst, D. et al. Characterization of the histone H2A.Z-1 and H2A.Z-2 isoforms in vertebrates. BMC Biol. 7, 1–16 (2009).

29. Schones, D. E. et al. Dynamic regulation of nucleosome positioning in the human genome. Cell 132, 887–898 (2008).

30. Chen, F. X., Smith, E. R. & Shilatifard, A. Born to run: control of transcription elongation by RNA polymerase II. Nat. Rev. Mol. Cell Biol. 19, 464–478 (2018).

31. Rahl, P. B. et al. C-Myc regulates transcriptional pause release. Cell 141, 432–445 (2010).

32. Cisse, I. I. et al. Real-Time Dynamics of RNA Polymerase II Clustering in Live Human Cells. Science 341, 664–667 (2013).

33. Steurer, B. et al. Live-cell analysis of endogenous GFP-RPB1 uncovers rapid turnover of initiating and promoter-paused RNA Polymerase II. Proc. Natl. Acad. Sci. 115, E4368–E4376 (2018).

34. Forero-Quintero, L. S. et al. Live-cell imaging reveals the spatiotemporal organization of endogenous RNA polymerase II phosphorylation at a single gene. bioRxiv 2020.04.03.024414 (2020). doi: 10.1101/2020.04.03.024414

35. Titov, D. V et al. XPB, a subunit of TFIIH, is a target of the natural product triptolide. Nat. Chem. Biol. 7, 182–188 (2011).

36. Gilchrist, D. A. et al. NELF-mediated stalling of Pol II can enhance gene expression by blocking promoter-proximal nucleosome assembly. Genes Dev. 22, 1921–1933 (2008).

37. Yung, T. M. C., Narita, T., Komori, T., Yamaguchi, Y. & Handa, H. Cellular dynamics of the negative transcription elongation factor NELF. Exp. Cell Res. 315, 1693–1705 (2009).

38. Fant CB, Levandowski CB, Gupta K, et al. TFIID Enables RNA Polymerase II Promoter-Proximal Pausing. Mol Cell. 78 (4):785–793 (2020).

39. Shao, W. & Zeitlinger, J. Paused RNA polymerase II inhibits new transcriptional initiation. Nat. Genet. 49(7):1045–1051 (2017).

40. Jonkers, I., Kwak, H. & Lis, J. T. Genome-wide dynamics of Pol II elongation and its interplay with promoter proximal pausing, chromatin, and exons. Elife 2014, 1–25 (2014).

41. Henriques, T. et al. Stable Pausing by RNA Polymerase II Provides an Opportunity to Target and Integrate Regulatory Signals. Mol. Cell 52, 517–528 (2013).

42. Kellner, W. A., Bell, J. S. K. & Vertino, P. M. GC skew defines distinct RNA polymerase pause sites in CpG island promoters. Genome Res. 25, 1600–1609 (2015).

43. Szlachta, K. et al. Alternative DNA secondary structure formation affects RNA polymerase II promoter-proximal pausing in human. Genome Biol. 19, 89 (2018).

44. Chen, F., Gao, X., Shilatifard, A. & Shilatifard, A. Stably paused genes revealed through inhibition of transcription initiation by the TFIIH inhibitor triptolide. Genes Dev. 29, 39–47 (2015).

45. Nogales, E., Louder, R. K. & He, Y. Cryo-EM in the study of challenging systems: the human transcription pre-initiation complex. Curr. Opin. Struct. Biol. 40, 120–127 (2016).

46. Zhang, Z. et al. Rapid dynamics of general transcription factor TFIIB binding during preinitiation complex assembly revealed by single-molecule analysis. Genes Dev. 30, 2106–2118 (2016).

47. de Graaf, P. et al. Chromatin interaction of TATA-binding protein is dynamically regulated in human cells. J. Cell Sci. 123, 2663–2671 (2010).

48. Danko, C. G. et al. Signaling pathways differentially affect RNA polymerase II initiation, pausing, and elongation rate in cells. Mol. Cell 50, 212–222 (2013).

49. Krebs, A. R. et al. Genome-wide Single-Molecule Footprinting Reveals High RNA Polymerase II Turnover at Paused Promoters. Mol. Cell 67, 411–422 (2017).

50. Mulas, C. et al. Defined conditions for propagation and manipulation of mouse embryonic stem cells. Development 146 (6) (2019).

51. Mylonas, C. & Tessarz, P. Transcriptional repression by FACT is linked to regulation of chromatin accessibility at the promoter of ES cells. Life Sci. Alliance 1 (3) (2018).

52. Langmead, B. & Salzberg, S. L. Fast gapped-read alignment with Bowtie 2. Nat Methods 9, 357–359 (2012).

53. Zhang, Y., Lin, Y. H., Johnson, T. D., Rozek, L. S. & Sartor, M. A. PePr: A peak-calling prioritization pipeline to identify consistent or differential peaks from replicated ChIP-Seq data. Bioinformatics 30, 2568–2575 (2014).

54. Ramirez, F. et al. deepTools2: a next generation web server for deep-sequencing data analysis. Nucleic Acids Res. 44, 160–165 (2016).

55. Hansen, P. et al. Q-nexus: a comprehensive and efficient analysis pipeline designed for ChIP-nexus. BMC Genomics 17, 873 (2016).

56. Love, M. I., Huber, W. & Anders, S. Moderated estimation of fold change and dispersion for RNA-seq data with DESeq2. Genome Biol. 15, 550 (2014).

57. Sorenson R, Johnson K, Adler F, S. L. RNAdecay: Maximum Likelihood Decay Modeling of RNA Degradation Data. (2019).

58. Sergé, A., Bertaux, N., Rigneault, H. & Marguet, D. Dynamic multiple-target tracing to probe spatiotemporal cartography of cell membranes. Nat. Methods 5, 687–694 (2008).

59. Andrews, J. O. et al. qSR: a quantitative super-resolution analysis tool reveals the cell-cycle dependent organization of RNA Polymerase I in live human cells. Sci. Rep. 8 (1), 7424 (2018).

60. Dobin, A. et al. STAR: Ultrafast universal RNA-seq aligner. Bioinformatics 29, 15–21 (2013).

61. Robinson, M. D., McCarthy, D. J. & Smyth, G. K. edgeR: A Bioconductor package for differential expression analysis of digital gene expression data. Bioinformatics 26, 139–140 (2009).

62. Ritchie, M. E. et al. limma powers differential expression analyses for RNA-sequencing and microarray studies. Nucleic Acids Res. 43 (7) e47 (2015).

63. Mayer, A. & Churchman, L. S. Genome-wide profiling of RNA polymerase transcription at nucleotide resolution in human cells with native elongating transcript sequencing. Nat Protoc. 11, 813–833 (2016).

64. Wamstad, J. A. et al. Dynamic and Coordinated Epigenetic Regulation of Developmental Transitions in the Cardiac Lineage. Cell 151, 206–220 (2012).

65. Illingworth, R. S. et al. The E3 ubiquitin ligase activity of RING1B is not essential for early mouse development. Genes Dev. 29, 1897–1902 (2015).

66. Mas, G. et al. Promoter bivalency favors an open chromatin architecture in embryonic stem cells. Nat. Genet. 50, 1452–1462 (2018).

