## Supplementary Figures for "A dual role for H2A.Z.1 in modulating the dynamics of RNA Polymerase II initiation and elongation"

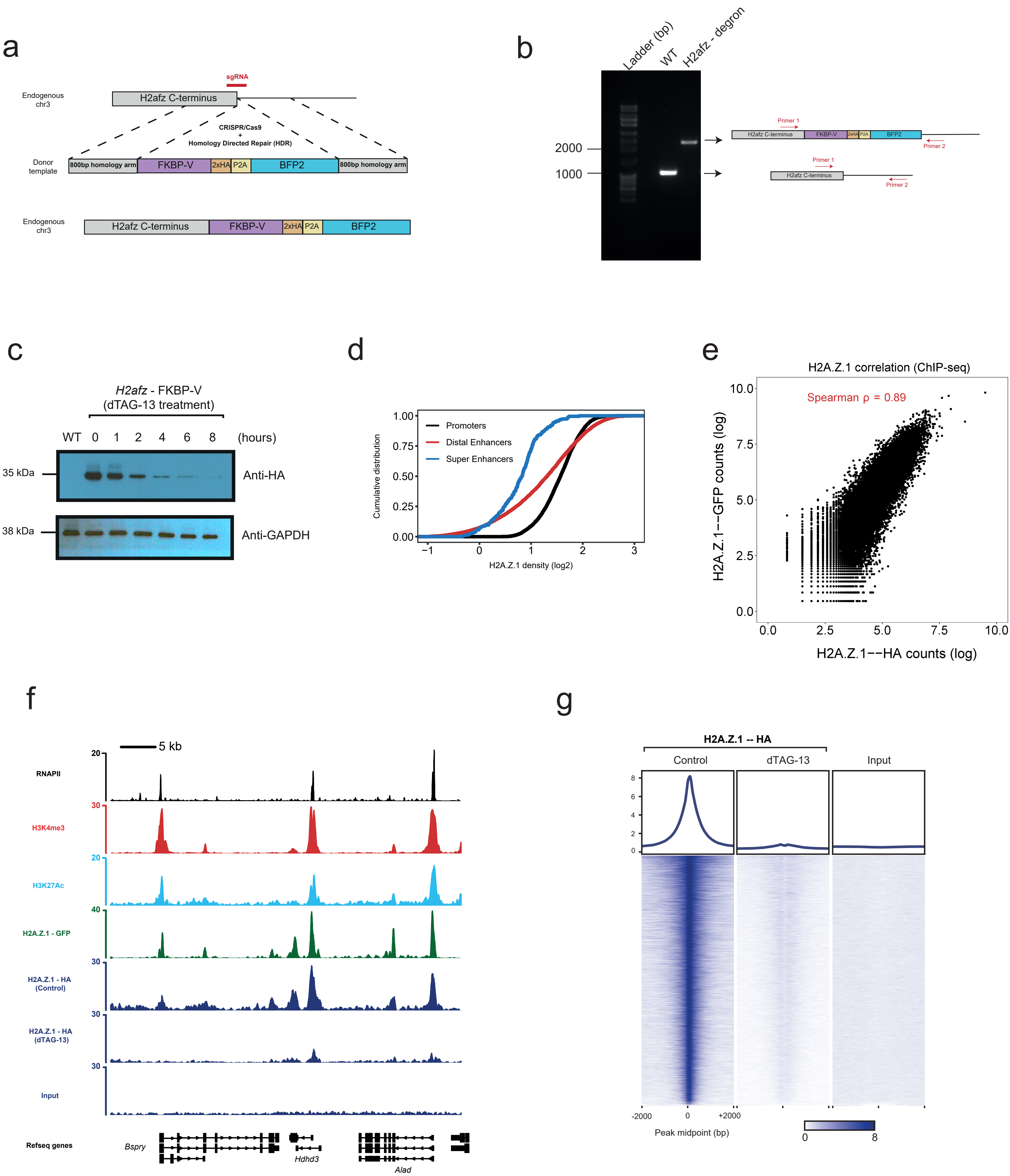


Extended Data Fig. 1

**Extended Data Fig.1 | Endogenous tagging of H2A.Z.1 with HA-FKBP-V in mESCs. a.** Schematic diagram of endogenous tagging of H2afz using CRISPR-Cas9 and homologous directed repair (HDR). **b.** Genotyping of WT and H2A.Z.1-FKBP-V mESCs cells using primers (Primer 1 & Primer 2) binding around the C-terminus of the *H2afz* gene. PCR amplified samples were run on a 1% agarose gel. **c.** Western blotting with Anti-HA and Anti-GAPDH antibodies in WT and H2A.Z.1-FKBP-V (Control or dTAG-13 treated) mESCs. H2A.Z.1-FKBP-V cells were treated with dTAG-13 for 0,1,2,4,6, and 8 hours. Maximum H2A.Z.1 depletion was observed after 8h of treatment with the small molecule. **d.** Cumulative distribution plot of H2A.Z.1 ChIP-seq over active protein-coding promoters (n = 7,789), active distal enhancers (n = 6,856), and super-enhancers (n=231). **e.** Scatterplot of H2A.Z.1-HA counts ($log_2$) and H2A.Z.1-GFP counts ($log_2$) over the promoters of 12,737 uniquely annotated protein-coding genes. Spearman correlation in indicated in red ($\rho = 0.89$). **f.** Single gene plots of several ChIP-seq datasets (RNAPII, H3K4me3, H3K27Ac, H2A.Z.1- GFP, H2A.Z.1-HA (Control & dTAG-13), Input). **g.** ChIP-seq heatmaps over 12,031 significant H2A.Z.1-HA peaks (FDR < $10^{-5}$) in Control and dTAG-13 treated cells. A heatmap of Input control is also presented.

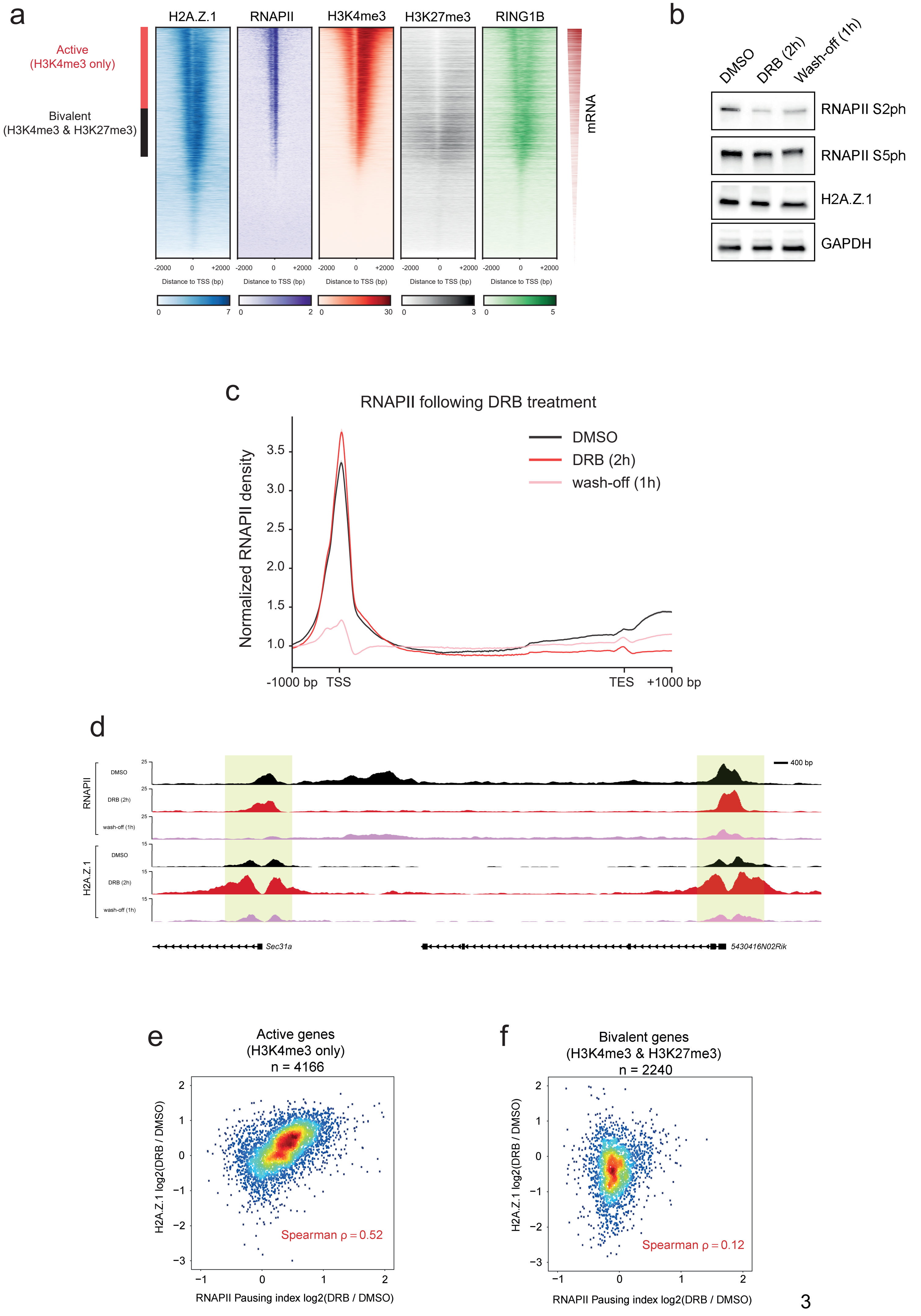




Extended Data Fig. 2

**Extended Data Fig.2 | H2A.Z.1 is a barrier to RNAPII progression. a.** ChIP-seq heatmaps over 10,878 uniquely annotated genes for H2A.Z.1, RNAPII, H3K4me3, H3K27me3, and RING1B. Genes are sorted by H3K4me3 levels. Corresponding mRNA levels are highlighted in red. **b.** Western blotting with Anti-Pol II S2ph, Anti-Pol II S5ph, Anti-HA (H2A.Z.1), and Anti-GAPDH after DRB treatment (2h) and subsequent wash-off (1h). DMSO concentration is at 2%. **v.** Average RNAPII metaplot profiles of 7,624 uniquely annotated genes for Control (DMSO), DRB, and wash-off over TSS, gene body, and TES. **d.** Single gene plots of ChIP-seq (H2A.Z.1 and RNAPII) for Control (DMSO), DRB, and wash-off conditions. **e.** Scatterplot of H2A.Z.1 ChIP-seq logFC (DRB / DMSO) versus RNAPII Pausing Index logFC (DRB / DMSO) over 4,166 Active genes (H3K4me3 only). Spearman correlation in indicated in red ($\rho$ = 0.52). **f.** Scatterplot of H2A.Z.1 ChIP-seq logFC (DRB / DMSO) versus RNAPII Pausing Index logFC (DRB / DMSO) over 2,240 Bivalent genes (H3K4me3 & H3K27me3). Spearman correlation in indicated in red ($\rho$ = 0.12).

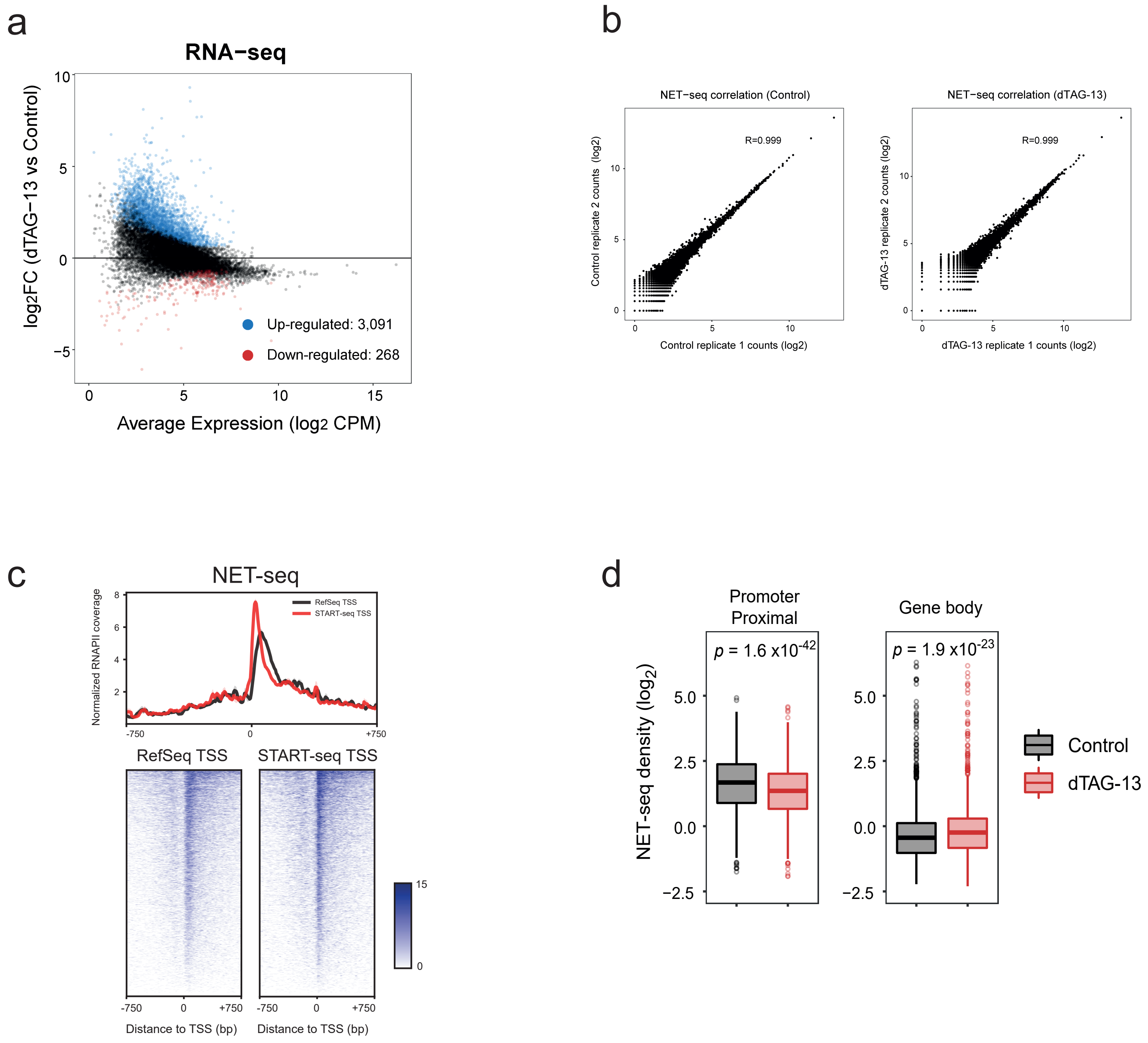


### Extended Data Fig. 3

**Extended Data Fig.3 | H2A.Z.1 acts as a transcriptional repressor. a.** MA plot of RNA-seq between dTAG-13 vs Control. Significant genes are highlighted in red and blue (logFC < -0.6 or logFC > 0.6 & adj.P.Value < 0.05). **b.** Metaplot profiles and heatmaps of RNAPII (NET-seq) over 4,184 protein-coding RefSeq TSS or START-seq TSS. **c.** Correlation plots for biological NET-seq replicates (Control – dTAG-13). **d.** Boxplots measuring either promoter proximal (-30 to +250 bp of TSS) or gene body (+300 bp to TES) RNAPII density between Control and dTAG-13 treated cells. Significance is calculated using a paired Wilcoxon rank test.

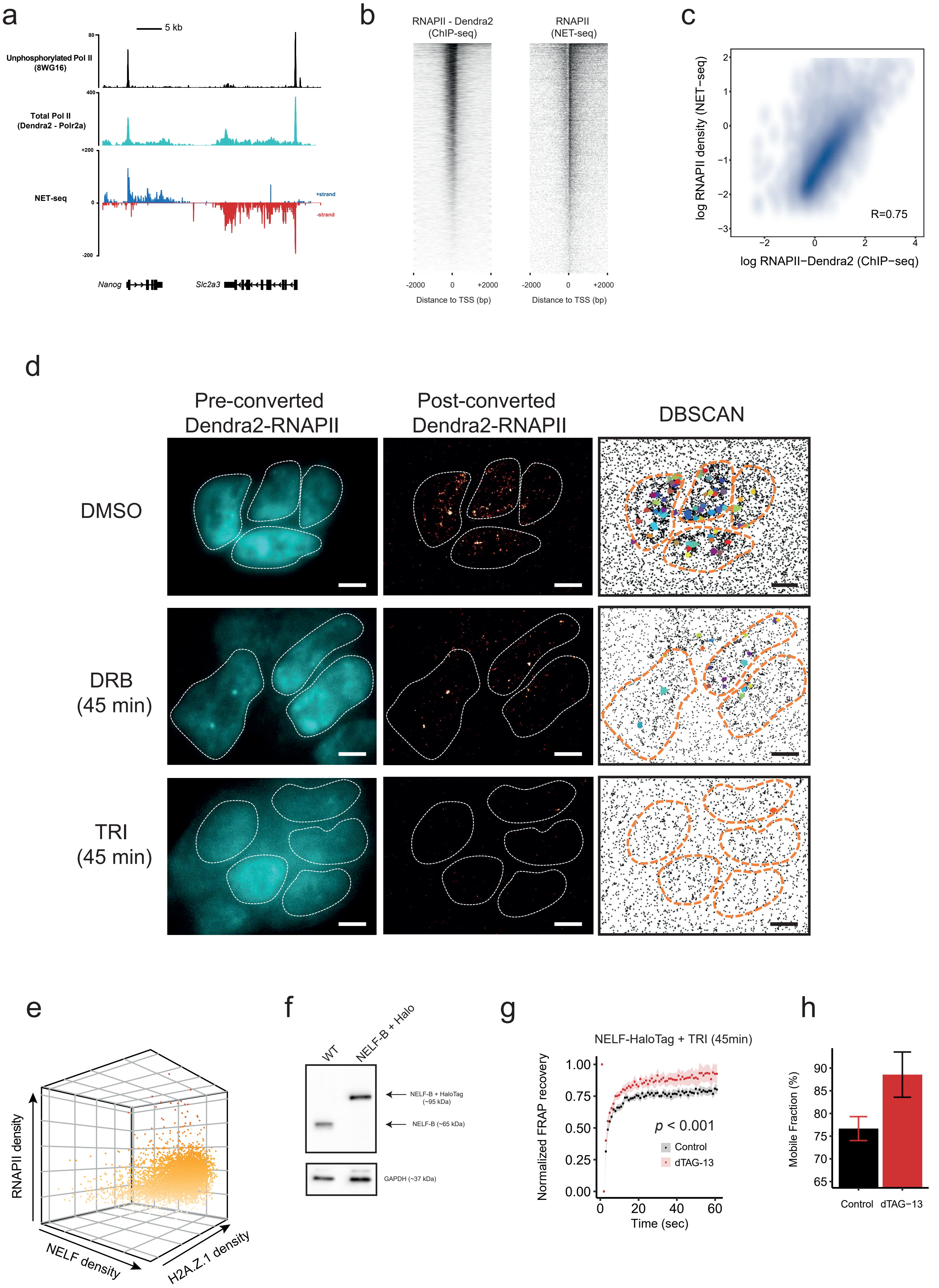


### Extended Data Fig. 4

**Extended Data Fig.4 | Rapid dynamics of endogenously tagged NELF-B and RNAPII. a.** Single gene plots of unphosphorylated RNAPII (ChIP-seq), Total RNAPII (Anti-Dendra2 ChIP-seq), and NET-seq. **b.** Heatmaps of Anti-Dendra2-RNAPII (ChIP-seq) and NET-seq over 11,315 protein-coding genes. **c.** Correlation plot between ChIP-seq (Anti-Dendra2-RNAPII) and NET-seq over 11,315 protein-coding genes. Pearson correlation is indicated on the plot (R = 0.75). **d.** Live-cell direct image of pre-converted Dendra2-RNAPII (left), super-resolution image of post-converted Dendra2-RNAPII (middle), and DBSCAN analysis of post-converted Dendra2-RNAPII (right). Cells were treated either with DMSO, DRB or TRI for 45 min before photo conversion. Scale bar 5 µm. **e.** 3D Scatterplot of RNAPII density, NELF density, and H2A.Z.1 density over 6,211 protein-coding genes. **f.** Western blotting with Anti-NELFB and Anti-GAPDH antibodies in WT and NELFB-Halo mESCs. **g.** The normalized recovery curve for NELF (*n* = 14 cells) in the presence of triptolide (TRI -45 min) yielded a recovery fraction of ~80% during the 60 sec observation in Control (black) conditions that increased to ~90% in dTAG-13 treated cells. Dots and shaded areas represent mean and SEM values, respectively. The Welch unpaired t-test was used for significance. **h.** Mobile fraction of in Control (black) and dTAG-13 (red) conditions plus TRI treatment. Error bars represent ±SEM.

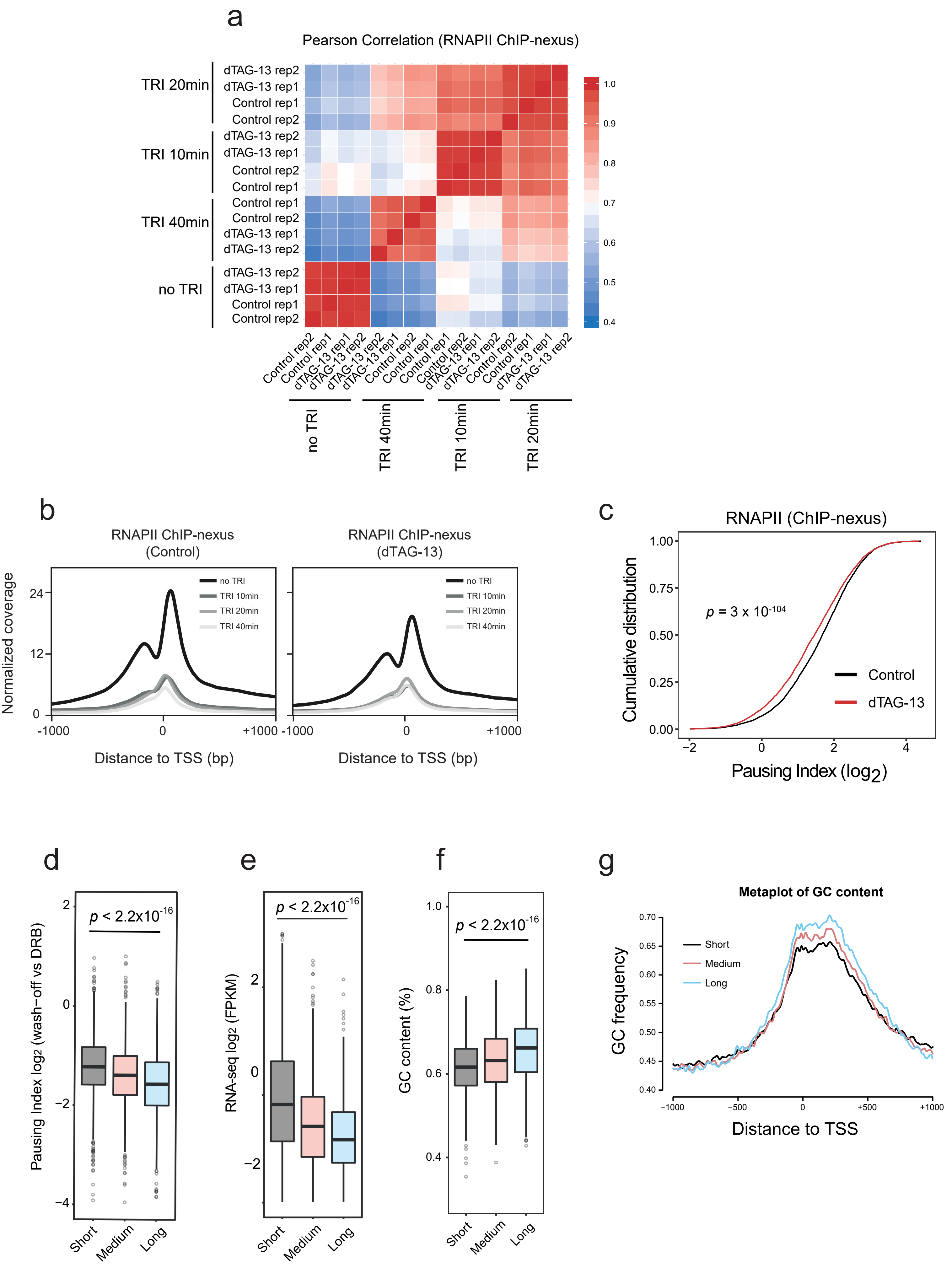


Extended Data Fig. 5

**Extended Data Fig.5 | RNAPII ChIP-nexus over the course of TRI treatment. a.** Pearson correlation heatmap of all RNAPII ChIP-nexus replicates (Control or dTAG-13 and no TRI, TRI 10 min, TRI 20 min, and TRI 40min). **b.** Metaplot profiles of RNAPII (ChIP-nexus) over 4,558 protein-coding TSSs for Control and dTAG-13 cells over the course of triptolide (TRI) treatment. **c.** Cumulative distribution plot of RNAPII (ChIP-nexus) Pausing Index for Control and H2A.Z.1-depleted (dTAG-13) mESCs, over 4,184 uniquely annotated protein-coding genes. Significance was calculated using a paired Wilcoxon rank test. **d.** Boxplots measuring $\log_2$ FC (wash-off / DRB) of RNAPII pausing index for the three different gene classes. Significance was calculated using an unpaired Wilcoxon rank test. **e.** Boxplots measuring mRNA abundance (FPKM) for the three different gene classes. Significance was calculated using an unpaired Wilcoxon rank test.**f.** Boxplots measuring GC content ± 250bp around the TSS for the three different gene classes. Significance was calculated using an unpaired Wilcoxon rank test. **g.** Metaplots of GC frequency for the three different gene classes ± 1000bp around the TSS.

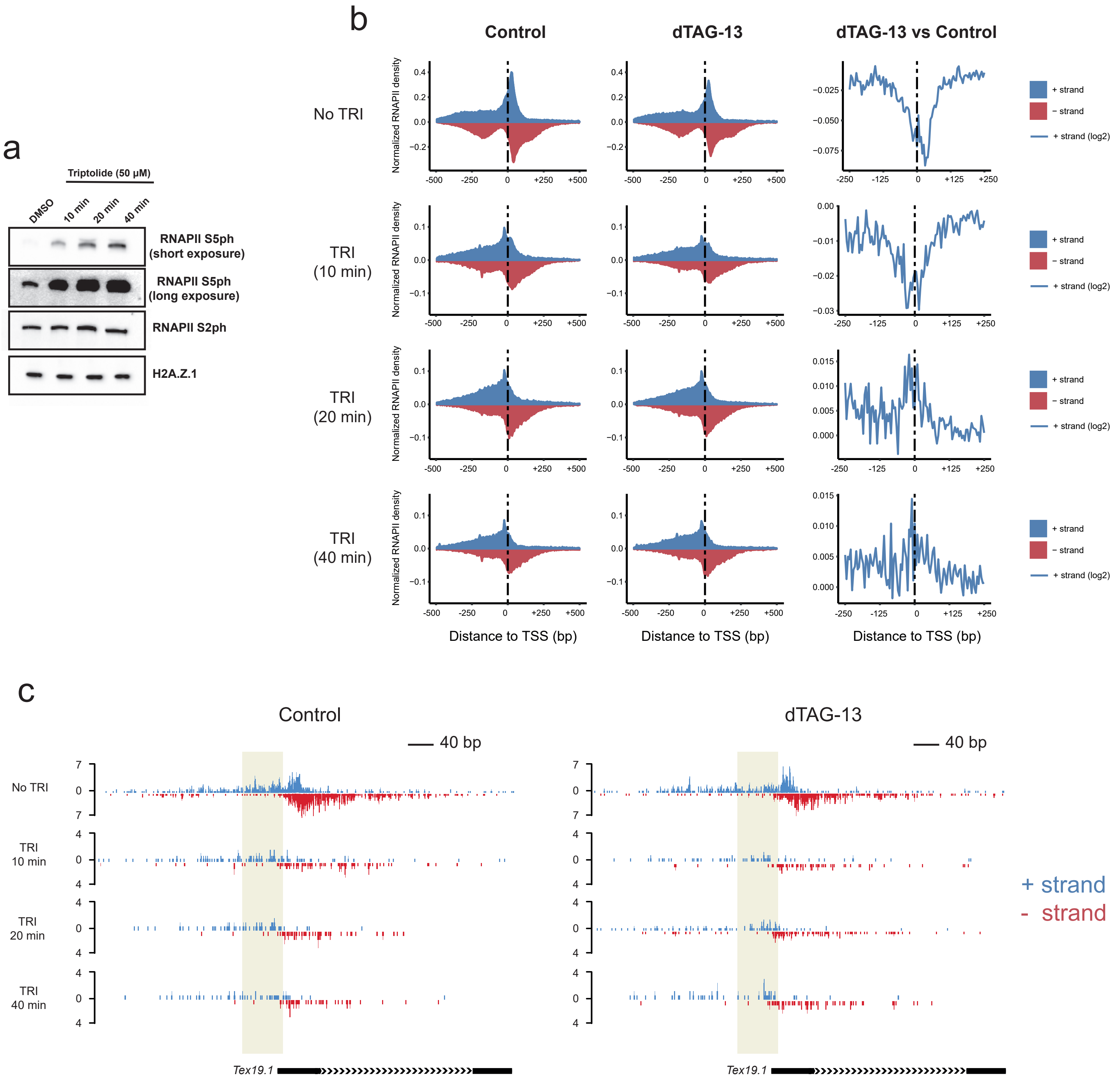


Extended Data Fig. 6

**Extended Data Fig.6 | TRI treatment reveals RNAPII at the PIC area. a.** Western blotting with Anti-Pol IIS5ph, Anti-Pol II S2ph, and Anti-HA (H2A.Z.1) antibodies over the course of TRI treatment. DMSO concentration is at 2%. **b.** Metaplot profiles of RNAPII (ChIP-nexus) over 1,460 protein-coding genes displaying the highest RNAPII at the PIC region. First column (Control), second column (dTAG-13), third column ($log_2$ dTAG-13/Control – Positive strand). **c.** Single gene plot (*Tex19.1*) of RNAPII ChIP-nexus profiles (Control, dTAG-13) over the course of Triptolide treatment (No TRI, 10 min, 20 min, and 40 min). RNAPII density is displayed both in the positive (blue) and negative (red) strand. Initiating RNAPII is highlighted in yellow.

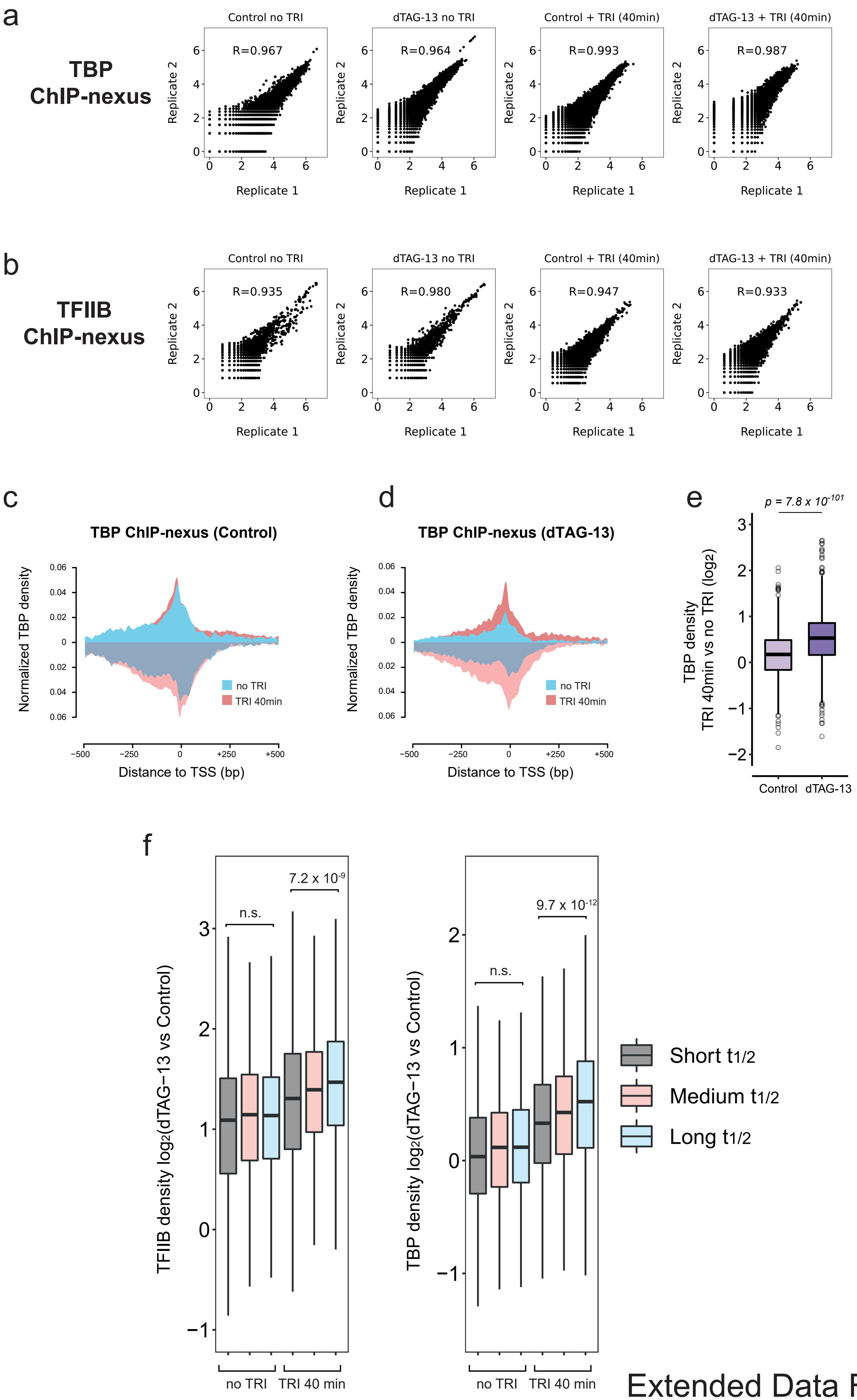




Extended Data Fig. 7

**Extended Data Fig.7 | TBP and TFIIB ChIP-nexus following TRI treatment. a.** Correlation plots for biological TBP ChIP-nexus replicates. **b.** Correlation plots for biological TFIIB ChIP-nexus replicates. **c.** Metaplots of TBP ChIP-nexus in Control conditions with (“TRI 40min”) or without (“No TRI”) triptolide. **d.** Metaplots of TBP ChIP-nexus in H2A.Z.1-depleted (dTAG-13) conditions with (“TRI 40min”) or without (“No TRI”) triptolide. **e.** Boxplots measuring $\log_2$ FC (TRI 40min vs No TRI) TBP density between Control and dTAG-13 in an area 250 bp upstream of the TSS (n = 2,143 genes). Significance was calculated using a Wilcoxon rank test. **f.** Boxplots measuring $\log_2$ FC (dTAG-13 / Control) of TFIIB and TBP at promoters of the three different gene classes. Significance was calculated using an unpaired Wilcoxon rank test.
